# MafF is an antiviral host factor that suppresses transcription from Hepatitis B Virus core promoter

**DOI:** 10.1101/2020.07.29.227793

**Authors:** Marwa K. Ibrahim, Tawfeek H. Abdelhafez, Junko S. Takeuchi, Kosho Wakae, Masaya Sugiyama, Masataka Tsuge, Masahiko Ito, Koichi Watashi, Mohamed El Kassas, Takanobu Kato, Asako Murayama, Tetsuro Suzuki, Kazuaki Chayama, Kunitada Shimotohno, Masamichi Muramatsu, Hussein H. Aly, Takaji Wakita

## Abstract

Hepatitis B Virus (HBV) is a stealth virus that exhibits only minimal induction of the interferon system that is required for both innate and adaptive immune responses. However, 90% of acutely infected adults can clear the virus, suggesting the presence of additional mechanisms that facilitate viral clearance. Herein, we report that Maf bZIP transcription factor F (MafF) promotes host defense against infection with HBV. Using siRNA library and an HBV/NL reporter virus, we screened to identify anti-HBV host factors. Our data showed that silencing of *MafF* led to a 6-fold increase in luciferase activity after HBV/NL infection. Overexpression of MafF reduced HBV core promoter transcriptional activity, which was relieved upon mutating the putative MafF binding region. Loss of MafF expression by CRISPR/CAS9 (in HepG2-hNTCP-C4 cells) or siRNA silencing (in primary hepatocytes [PXB]), induced HBV core and HBV pregenomic RNA (pgRNA) levels, respectively, after HBV infection. MafF physically binds to HBV core promoter and competitively inhibits HNF-4α binding to an overlapping sequence in HBV enhancer II sequence (EnhII) as seen by ChIP analysis. MafF expression was induced by IL-1β/TNF-α treatment in both HepG2 and PXB cells, in an NF-κB-dependent manner. Consistently, *MafF* expression levels were significantly enhanced and positively correlated with the levels of these cytokines in patients with chronic HBV infection, especially in the immune clearance phase.

**Importance:** HBV is a leading cause of chronic liver diseases, infecting about 250 million people worldwide. HBV has developed strategies to escape interferon-dependent innate immune responses. Hence, the identification of other anti-HBV mechanisms is important for understanding HBV pathogenesis, and developing anti-HBV strategies. MafF was shown to suppress transcription from HBV core promoter, leading to a significant suppression of HBV life cycle. Furthermore, MafF expression was induced in chronic HBV patients and in primary human hepatocytes (PXB). This induction correlated with the levels of inflammatory cytokines (IL-1β and TNF-α). These data suggest that the induction of MafF contributes to the host’s antiviral defense by suppressing transcription from selected viral promoters. Our data shed light on a novel role for MafF as anti-HBV host restriction factor.

## Introduction

In the earliest stages of viral infection, the host initially detects and counteracts infection via induction of innate immune responses (46). Host restriction factors are essential components of the innate antiviral immune response; these factors serve critical roles in limiting virus replication before the adaptive immune response engages to promote virus clearance (5). These antiviral restriction factors are typically induced by cytokines, including interferons (IFNs) (56), transforming growth factor-beta (TGF-β) (29), and interleukin-1-beta (IL-1β) (58). These restriction factors suppress viral replication by targeting the infection at various stages of the virus life cycle, including viral entry (3), transcription of the viral genome (67), viral RNA stability (2), translation of viral proteins (33), viral DNA replication (34), and production of viral particles (6).

Approximately 250 million people worldwide are chronically infected with Hepatitis B virus (HBV). These patients are at high risk of developing life-threatening complications, including hepatic cirrhosis, hepatic failure, and hepatocellular carcinoma. Current treatments include nucleos(t)ide analogs that efficiently suppress HBV replication. However, an HBV replication intermediate, covalently closed circular DNA (cccDNA), persists in the nucleus. The cccDNA intermediate gives rise to progeny virus, and may lead to the development of drug-resistant mutants and/or relapsing HBV after drug withdrawal (43). As such, new strategies for HBV treatment are needed.

HBV has been identified in human remains from ∼7000 years ago (27). This prolonged history and evolution has shaped HBV to be one of the most successful of the “stealth” viruses that can successfully establish infection while evading IFN induction (66). Although HBV can evade IFN induction, the majority of HBV-infected adults (90%) are ultimately able to clear the virus. This observation suggests that there are likely to be one or more IFN-independent host restriction factors that facilitate HBV clearance.

The small Maf proteins (sMafs) are a family of basic-region leucine zipper (bZIP)-type transcription factors. MafF, MafG and MafK are the three sMafs identified in vertebrate species (24). sMafs lack a transcriptional activation domain hence, they can act as both transcription activator or repressor based on their expression levels and dimerization partners (19). Intriguingly, previous reports have documented induction of MafF in myometrial cells by inflammatory cytokines, including IL-1β and tumor necrosis factor alpha (TNF-α) (32). However, there have been no previous studies that have addressed a role for MafF in promoting an antiviral innate immune response.

Using an HBV reporter virus and an siRNA library, we performed functional siRNA screening to identify the host factors that influence the HBV life cycle. Based on the results of this screen, we identified MafF as a negative regulator of HBV infection. Further analysis revealed that MafF functions as a repressor of transcription at the HBV core promoter, thereby suppressing HBV replication. This is the first study to report a role for MafF as an anti-HBV host factor that represses transcription from the promoters of susceptible viruses.

## Results

### 1. MafF suppresses expression of the HBV/NanoLuc (NL) reporter virus

HBV particles carrying a chimeric HBV virus encoding NanoLuc (NL) were prepared as previously described (45). Since these particles carry a chimeric HBV genome in which HBc is replaced by NL, the NL levels released after infection with these particles can only detect the early stages of HBV infection from entry to transcription of HBV-pgRNA (45). We used this high-throughput system, in combination with druggable genome siRNA Library, to screen for host factors that influence these early stages. This approach facilitated testing of 2200 human genes for their influence on the HBV life cycle. Screening was performed in HepG2-C4 cells that express the HBV entry receptor, hNTCP (17). Non-targeting siRNAs, and siRNAs against h*NTCP*, were used as controls for each plate (Fig. 1A). Cellular viability was determined using the XTT assay; wells with ≥20% loss of cell viability were excluded from further evaluation. NL activity was induced more than 5 folds (average of 3 different siRNAs) upon the independent silencing of only 10 out of the 2200 host genes (0.4%). These genes were identified as anti-HBV host factors, and based on the induction level of NL activity, these genes were classified into 3 groups: Genes in which NL activity ranged from 5 to 10 folds (n=6 genes, MafF fits in this group), from 10 to 20 folds (3 genes), and from 20 to 30 folds induction of NL activity (1 gene) (Fig. 1A). MafF was one of the anti-HBV host factors identified by this screening. MafF was previously reported to be induced by famous inflammatory cytokines (IL-1β, and TNF-α), one of the common criteria of anti-viral host restriction factors (29, 56, 58), hence we decided to analyze its role on HBV life cycle. Silencing of *MafF* expression with si-1 or si-3 resulted in 6- (*p* < 0.0001) or 10-fold (*p* < 0.001) increases in NL activity, respectively, compared to that observed in cells transfected with the control siRNA (Fig. 1B). The MafF-specific sequence, si-2, did not show a similar effect on NL activity (Fig. 1B). This result was consistent with the fact that si-2 had a lower silencing efficiency for MafF (Fig. 1C). Since NL activity was measured 10 days after siRNA silencing of *MafF* expression, we measured the MafF protein levels at the designated time in order to confirm the prolonged silencing of *MafF* by si-3 (Fig. 1D). Taken together, these findings suggest that MafF may suppress HBV infection.

**Figure 1.**
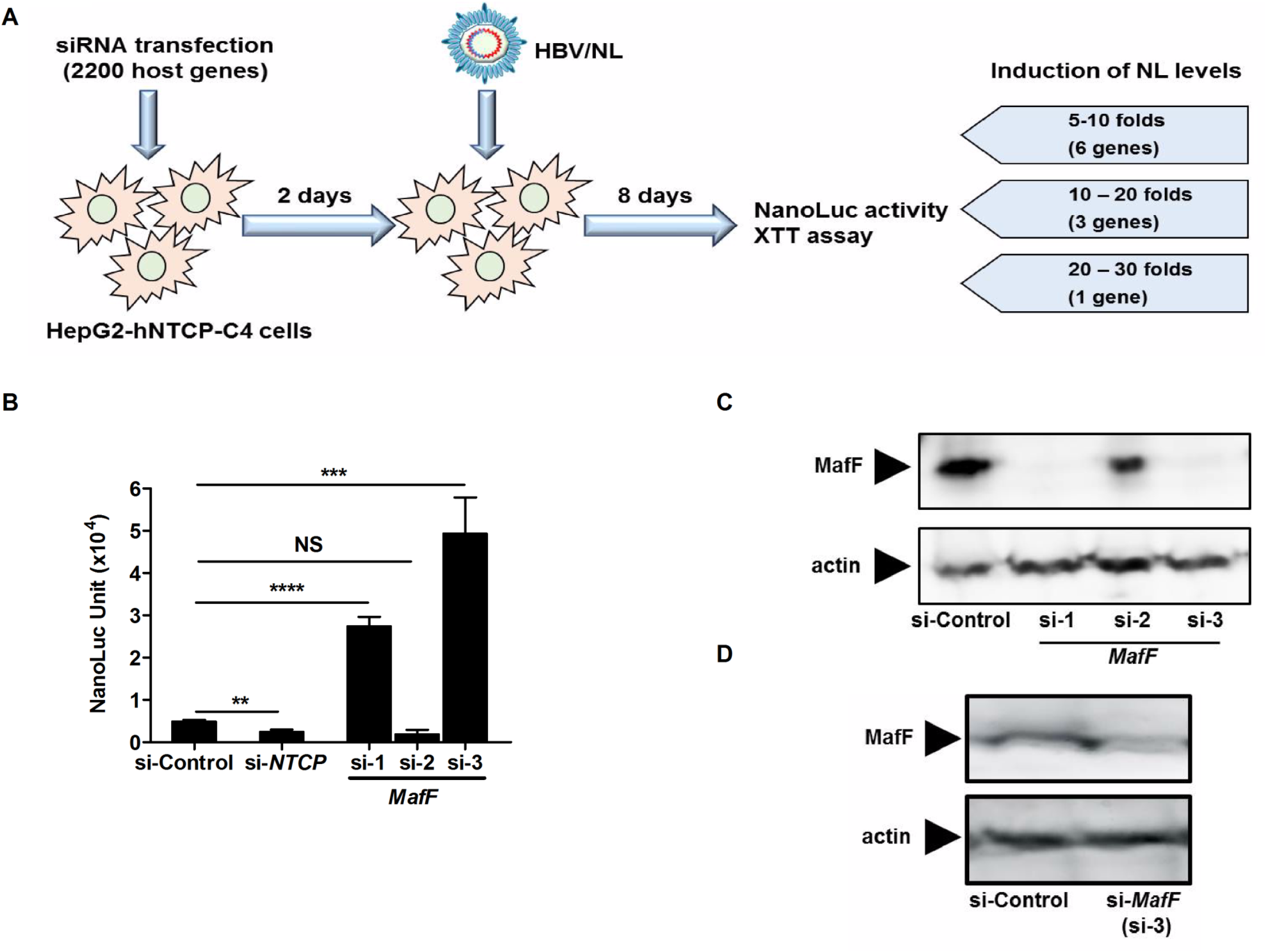
MafF suppresses HBV infection. **A.** A schematic diagram showing the experimental approach used to screen the siRNA library. **B.** HepG2-hNTCP-C4 cells were transfected with control, *NTCP*, or *MafF*-targeting siRNAs (si-1, si-2, and si-3); two days later, transfected cells were infected with the HBV/NL reporter virus. At day 8 post-infection, luciferase assays were performed, and NanoLuc activity was measured and plotted. **C.** HepG2 cells were transfected with control or *MafF*-targeting siRNAs (si-1, si-2, and si-3); total protein was extracted after two days. Expression of MafF (upper panel) and actin (control; lower panel) was analyzed by immunoblotting with their respective antibodies. **D.** HepG2 cells were transfected with control or *MafF*-targeting siRNA (si-3); total protein was extracted after ten days. Expression of MafF (upper panel) and actin (control; lower panel) was analyzed by immunoblotting with their respective antibodies. All assays were performed in triplicate and included three independent experiments. Data are presented as mean±standard deviation (SD); \*\**p*<0.01, \*\*\**p*<0.001, \*\*\*\**p*< 0.0001; NS, not significant

### 2. MafF strongly suppresses HBV core promoter activity

The HBV/NL reporter system can be used to detect factors affecting the early steps of the HBV life cycle, from HBV entry through cccDNA formation, transcription and translation of HBV-pgRNA (45). Silencing of *MafF* had no impact on cccDNA levels observed in cells infected with HBV as shown by real-time PCR (Fig. 2A) and by southern blot (Fig. 2B); these results indicated that MafF suppressed the HBV life cycle at stage that was later than that of cccDNA formation. Given that MafF can induce transcriptional suppression (19), we analyzed the impact of MafF on various HBV promoters (core, X, preS1, and preS2) using a reporter system in which firefly luciferase coding sequence was inserted downstream to the corresponding HBV promoter. We found that overexpression of MafF resulted in significant suppression of transcription from the HBV core promoter (approximately 8-fold; *p*<0.0001), and significant, albeit less of an impact on transcription from the HBV-X and preS1 promoters (both at approximately 2-fold, *p*<0.0001); overexpression of MafF had no significant impact on transcription from the preS2 promoter (Fig. 2C left panel). Likewise, siRNA silencing of endogenous *MafF* enhanced HBV core promoter activity (Fig. 2C right panel, *p*<0.0001). Since the NanoLuc gene in HBV/NL virus (Fig. 1B) is transcribed from an HBV core promoter (45), the findings presented in Fig. 1 and Fig. 2 collectively suggest that MafF-mediated suppression of HBV is mediated primarily by inhibition of transcription from the core promoter.

**Figure 2.**
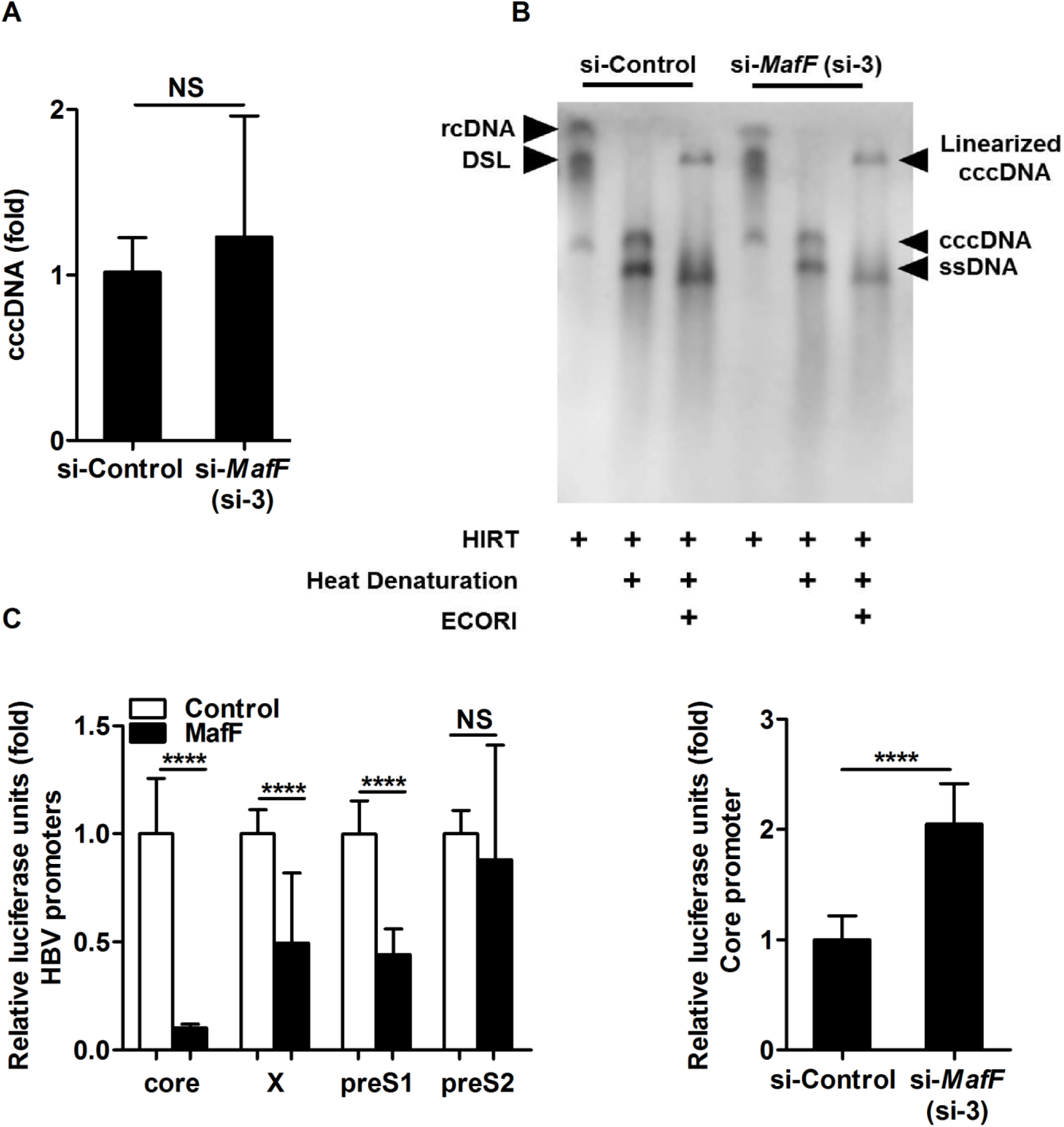
MafF suppresses the transcriptional activity of the HBV core promoter. **A.** HepG2-hNTCP-C4 cells were transfected with control or *MafF*-targeting siRNA (si-3). Two days after transfection, the transfected cells were infected with HBV at 12,000 genomic equivalents (GEq) per cell. Eight days later, the cells were harvested, DNA was extracted, and cccDNA was quantified by real-time PCR. The data were normalized to the levels of endogenous *GAPDH DNA* and are presented as fold change relative to control siRNA-transfected cells. **B.** HepG2-hNTCP-C4 cells were transfected with control or *MafF*-targeting siRNA (si-3). Two days after transfection, the transfected cells were infected with HBV at 12,000 GEq per cell. Eight days later, the cells were harvested, HIRT purification of DNA was performed, and cccDNA was visualized by southern blotting assay. **C.** HepG2 cells (left panel) were co-transfected with a MafF expression vector or empty vector (control) together with firefly luciferase reporter plasmids with HBV promoters (core, X, S1, and S2) and the pRL-TK control plasmid encoding *Renilla* luciferase. Two days after transfection, the cells were harvested and evaluated by dual luciferase assay. HepG2 cells (right panel) were transfected with control or *MafF*-targeting siRNA (si-3); 24 hours later, the cells were transfected with firefly luciferase reporter-HBV core promoter vector and the pRL-TK plasmid encoding *Renilla* luciferase. Two days later, the cells were lysed and evaluated by dual luciferase assay. For panel C, firefly luciferase data were normalized to *Renilla* luciferase levels; relative light units (RLUs) for firefly luciferase were plotted as fold differences relative to the levels detected in the control groups. All assays were performed in triplicate and included three independent experiments; data are presented as mean±SD. \*\*\*\**p*<0.0001; NS, not significant.

### 3. MafF suppresses HBV replication

The HBV core promoter controls the transcription of the longest two HBV RNA transcripts, the precore and pgRNAs. HBeAg is translated from the HBV precore RNA, while translation of HBV-pgRNA generates both the polymerase (Pol) and the capsid subunit; the pgRNA also serves as the template for HBV-DNA reverse transcription (4, 14). As such, we assumed that MafF served to inhibit HBV replication by controlling transcription of the HBV core promoter. In fact, overexpression of MafF resulted in significant suppression of the pgRNA titer of HBV genotypes A (GenBank: AB246338.1) and D (GenBank: V01460.1), as demonstrated by RT-qPCR (Fig. 3A, *p*<0.0001 for each genotype). Overexpression of MafF also suppressed the release of HBeAg as measured by enzyme-linked immunosorbent assay (ELISA) (Fig. 3B, *p*<0.0001), as well as the intracellular accumulation of HBV core protein as detected by immunoblotting (Fig. 3C upper and lower panels; *p*<0.05 by densitometric analysis) and the level of HBV core-associated DNA as revealed by southern blot (Fig. 3D).

**Figure 3.**
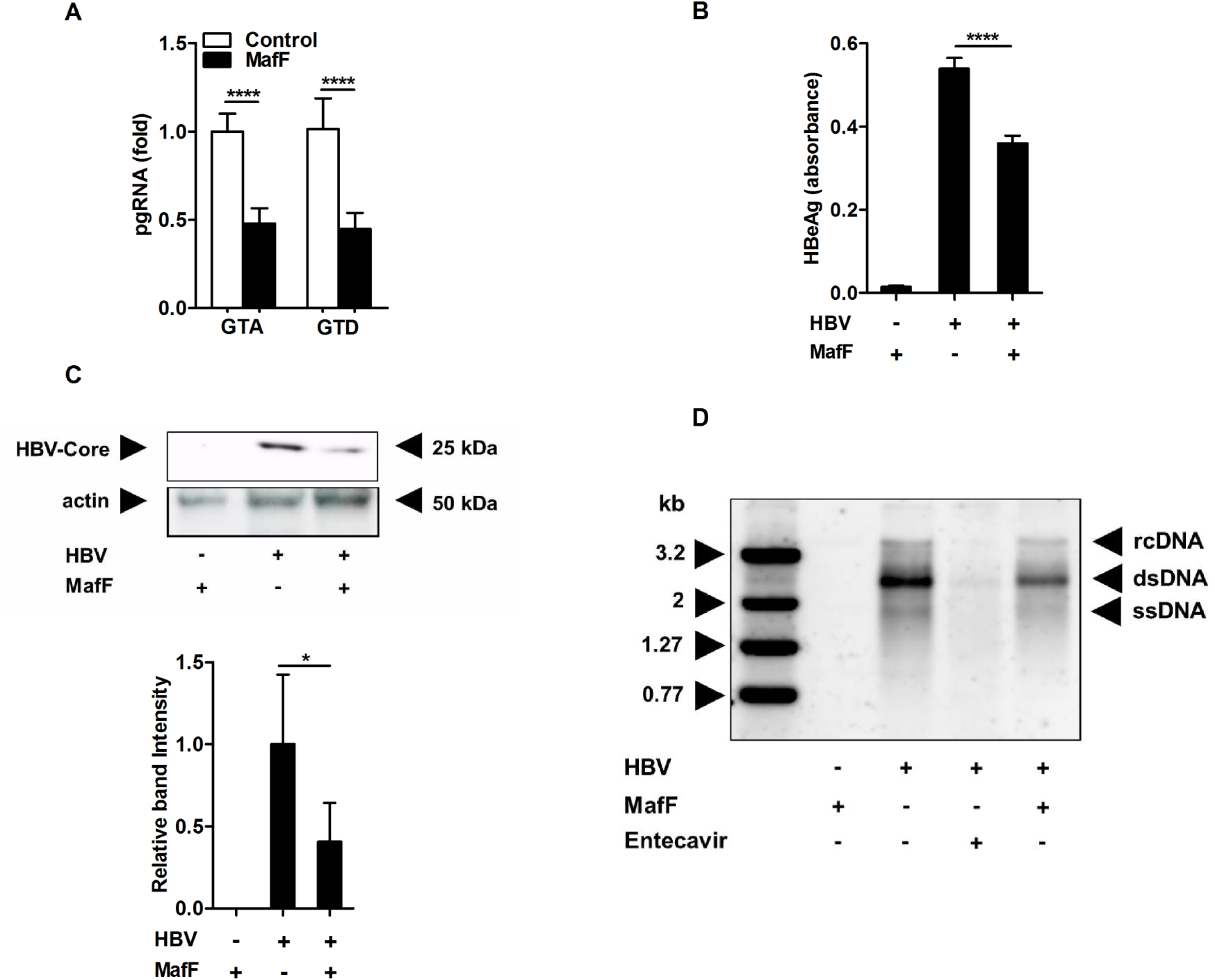
MafF suppresses HBV life cycle. **A.** HepG2 cells were transfected with empty (control) or MafF expression vector together with expression vectors encoding HBV genotypes A and D. Two days later, the cells were harvested and the pgRNA expression was quantified by real-time RT-PCR. The data were normalized to the expression of *GAPDH* and are shown as the fold change relative to control plasmid-transfected cells. **B–D:** HepG2 cells were transfected with empty (control) or MafF expression vectors together with an expression plasmid encoding HBV genotype D **(B)** At 2 days post-transfection, HBeAg in the cell culture supernatants were quantified by ELISA. **(C)** The intracellular levels of HBV core protein (upper left panel) and actin (loading control; lower left panel) were evaluated by immunoblotting; the intensities of the bands (right panel) were quantified by ImageJ software. **(D)** At 3 days post-transfection, the levels of intracellular core-associated DNA were determined by southern analysis; transfected cells treated with 10 µM entecavir were used as controls. Data are presented as fold differences relative to the control plasmid-transfected cells. All assays were performed in triplicate and include results from three independent experiments; data are presented as mean±SD; \**p*<0.05, \*\*\*\**p*<0.0001.

### 4. MafF-KO induce HBV core protein levels

To further clarify the significance of MafF on HBV infection, we established CRISP/CAS9 MafF-KO HepG2-hNTCP-C4 cells. Out of 11 selected clones, MafF-KO-8 and 11 showed the best KO phenotype (Fig. 4A). Myrcludex-B is a lipopeptide consisting of amino acid residues 2–48 of the pre-S1 region of HBV, and is known to block HBV entry [18], pre-treatment with Myrcludex-B (1 μM) 1 hour before infection was performed to confirm that the detected signals were derived from HBV infection and did not represent non-specific background(12). Both MafF-KO-8 and 11 showed a higher NL secretion after HBV/NL infection when compared to parental HepG2-hNTCP-C4 cells with values ranging from 1.5 to 3 folds respectively (Fig. 4B). Accordingly, MafF-KO-11 showed 4 times higher HBc levels after HBV infection (Fig. 4C right and left panels). In comparison to the original HepG2-hNTCP-C4 cells, MafF-KO-11 cells showed similar levels of secreted HBs after HBV infection (Fig. 4D), which can be explained by the major function of MafF as a transcriptional repressor of HBV core promoter with minimal to no effect of PreS1 and PreS2 promoters (Fig. 2 B and C).

**Fig. 4.**
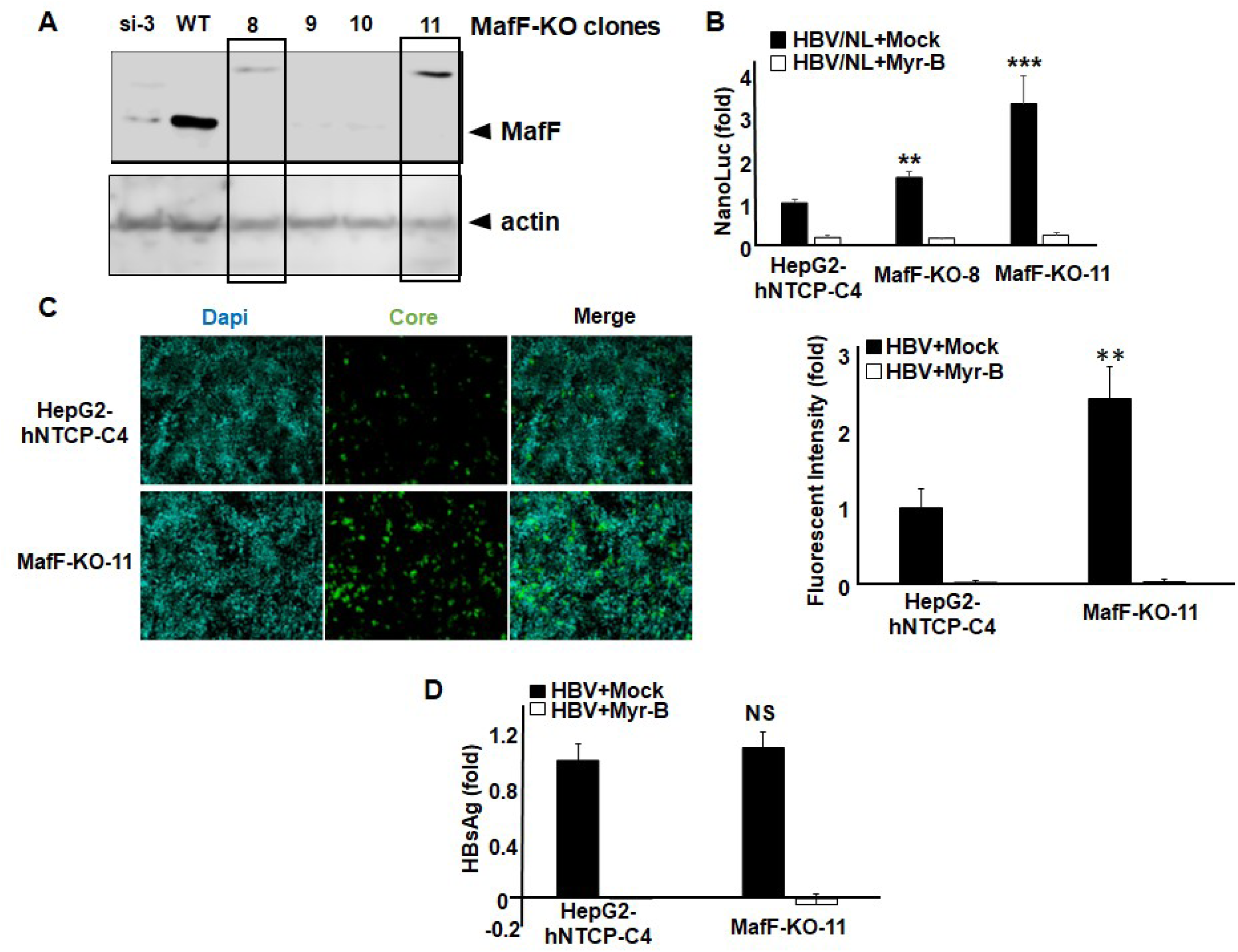
Enhanced expression of HBc in MafF-KO cells. A. HepG2-hNTCP-C4 cells were co-transfected with MAfF CRISPR/Cas9 KO Plasmid (h), sc-411785, and MAfF HDR Plasmid (h), sc-411785-HDR. Puromycin selection was conducted at 3μg/mL for 2 weeks. 11 isolated colonies were picked and scaled up. MafF expression was detected by immunoblotting, as positive and negative controls, we used WT (HepG2-hNTCP-C4) cells, and si-3 treated HepG2-hNTCP-C4 cells respectively. B. HepG2-hNTCP-C4, MafF-KO-8, and MafF-KO-11 cells were infected with HBV/NL reporter virus. Cells were pretreated with or without 1 μM Myrcludex-B or vehicle (DMSO) for 3 h before infection. At day 8 post-infection, luciferase assays were performed, and NanoLuc activity was measured and plotted as fold-difference relative to the mean luciferase levels in HepG2-hNTCP-C4 cells. C. HepG2-hNTCP-C4, or MafF-KO-11 cells were infected with HBV (6000 GEq/cell), at 12 days post-infection, HBc in the cells was detected by immunofluorescent (left panel) and fluorescent intensity was plotted as fold-difference relative to its mean levels in HBV-infected HepG2-hNTCP-C4 cells. D. HepG2-hNTCP-C4, or MafF-KO-11 cells were infected with HBV (6000 GEq/cell). Cells were pretreated with or without 1 μM Myrcludex-B or vehicle (DMSO) for 3 h before infection. After 12 days from infection, HBsAg in the cell culture supernatants was measured and plotted as fold-difference relative to the its mean levels in the supernatant of HBV-infected HepG2-hNTCP-C4 cells. All assays were performed in triplicate and include results from three independent experiments; data are presented as mean±SD; \*\**p*<0.01, \*\*\**p*<0.001.

### 5. MafF binds to the HBV core promoter

MafF is a member of (Maf) family of transcription factors, bZIP-type transcription factors that bind to DNA at Maf recognition elements (MAREs). MAREs were initially defined as a 13-bp (TGCTGA(G/C)TCAGCA) or 14-bp (TGCTGA(GC/CG)TCAGCA) elements (22, 25). The specificity of this binding sequence is greatly affected by the dimerization partners of MafF; multiple studies have presented findings suggesting heterogeneity within MARE sequences, especially when MafF heterodimerizes with anotherbZIP-type transcription factors (26, 44, 53). We next analyzed the HBV core promoter for putative MafF binding region using the JASPAR database of transcription factor binding sites (55). Toward this end, we identified the sequence 5’-TGGACTCTCAGCG-3’ that corresponded to nucleotides (nts) 1667 to 1679 of the HBV-C_JPNAT genome (GenBank AB246345.1) in the enhancer 2 (EnhII) of the HBV core promoter. This motif shared a similarity to a previously defined Maf responsive element (MARE) (19) and also with the DNA binding site for other cap’n’collar (CNC) family proteins Nrf1, Nrf2, Bach1 (Jaspar matrix profiles, MA0506.1, MA0150.1, and MA0591.1, respectively), andbZIP transcription factors that are reported to heterodimerize with sMafs (Fig. 5A). As such, we evaluated the role of this predicted MafF/bZIP site with respect to HBV core promoter activity. We found that the 9^th^ and 11^th^ nucleotides of the aforementioned predicted MafF binding region, which are A and C, respectively, are highly conserved common residues in the predicted MafF/bZIP binding sequence (Fig. 5A). We disrupted this predicted MafF/bZIP site by introducing 2-point mutations (A1676C and C1678A) into the HBV core promoter (Fig. 5A). Despite a minimal but statistically significant reduction of HBV core promoter activity induced by the introduction of these mutations (Fig. 5B left panel, *p*<0.0001), MafF overexpression suppressed the wild-type (WT) core promoter 2–3 times more than that carrying mutations (A1676C and C1678A) (Fig. 5B right panel, *p*<0.0001). Furthermore, ChIP analysis revealed that there was significantly less physical interaction between MafF and the HBV core promoter with A1676C and C1678A mutations than was observed between MafF and the HBV WT counterparts (Fig. 5C, *p*<0.05 for % of input and *p*<0.01 for fold enrichment). These results confirmed that MafF physically binds to the WT HBV core promoter at the putative MafF/bZIP binding region and thereby suppresses transcription. Interestingly the 5’ end of the identified MafF/bZIP binding region in HBV core promoter showed high conservation in all HBV genotypes, including ancient HBVs from the Bronze age to the early modern age (appx. 4,500 - 250 years ago), while 3’ end showed a relatively less conservation in HBV genotypes A to E, with more sequence divergence in new world HBV genotypes F, G and H (Fig. 5D). The role of these mutations in the escape from MafF-mediated transcriptional repression in these genotypes needs to be further analyzed.

**Figure 5.**
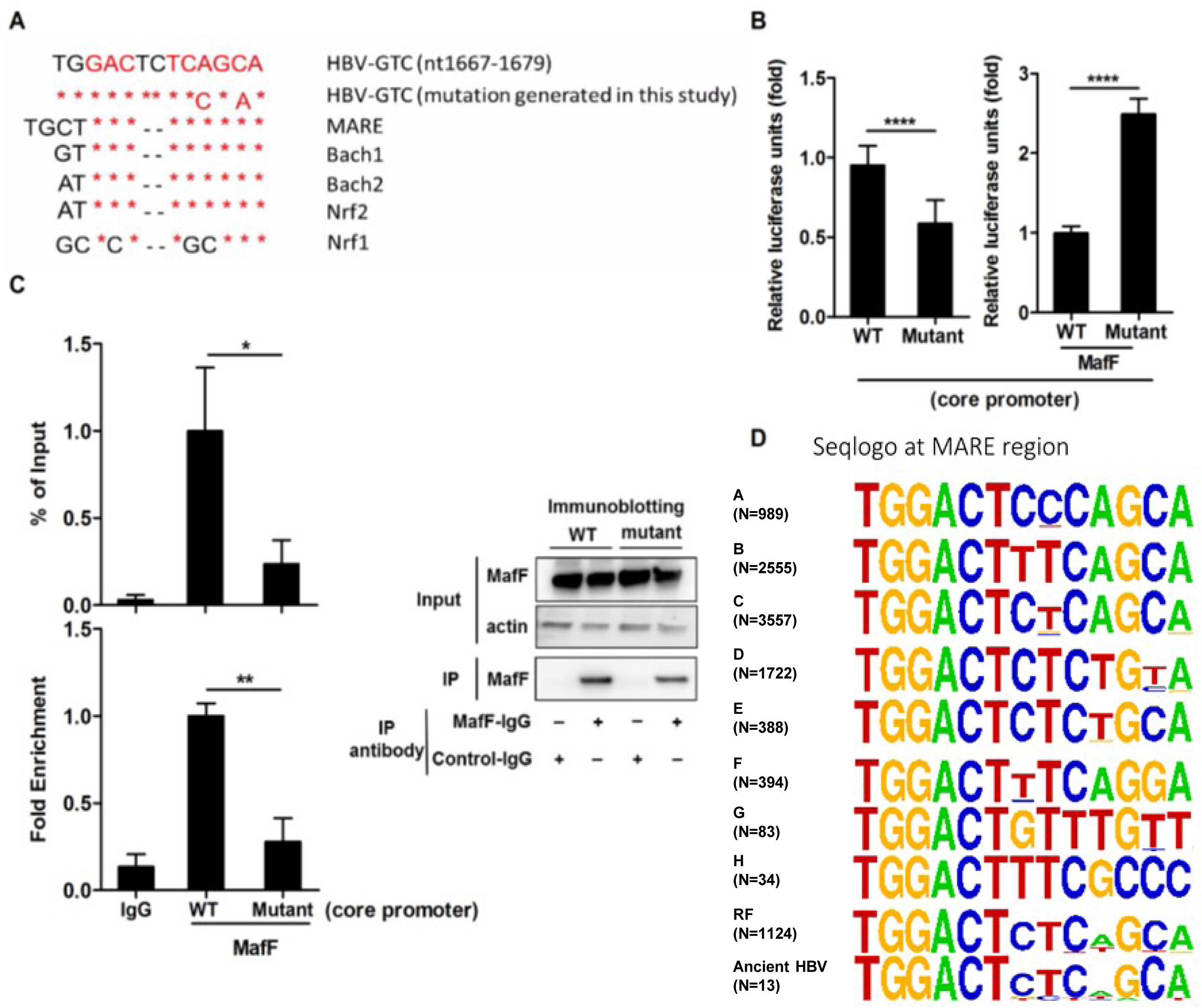
Physical interaction of MafF with HBV core promoter is required for transcriptional repression. **A.** A schematic representation of the putative MafF/bZIP binding region within enhancer 2 (EnhII) of the HBV core promoter from HBV genotype C. A mutant construct was prepared by introducing two point mutations (A1676C and C1678A) into the MARE sequence identified in the wild-type (WT) core promoter. **B.** HepG2 cells were co-transfected with mock (left panel) or a MafF expression plasmid (right panel) along with an HBV core promoter (WT or mutant)-reporter plasmid and pRL-TK encoding *Renilla* luciferase. At two days post-transfection, a dual luciferase assay was performed; firefly luciferase data were normalized relative to *Renilla* luciferase levels, and RLUs for firefly luciferase are plotted as fold differences relative to activity in the control group. **C.** 293FT cells were transfected with either the WT or mutant HBV core promoter-luciferase reporter plasmid together with a MafF expression plasmid (at a ratio 1:4). At two days post-transfection, cell lysates were collected; two aliquots (1/10 volume each) were removed from each sample. One aliquot was used for the detection of MafF protein (Input) and actin (loading control) by immunoblotting (right upper and middle panels); the second aliquot was used for DNA extraction and detection of HBV core promoter (Input) by real-time PCR. The remaining cell lysates (each 8/10 of the original volume) were subjected to ChIP assay using either isotype control antibody (rabbit IgG) or rabbit anti-MafF IgG to detect MafF. Following immunoprecipitation (IP), 1/10 volume of each IP sample was analyzed by immunoblotting for MafF (right lower panel); each remaining IP sample was subjected to DNA extraction and real-time PCR assay in order to detect associated HBV core promoter DNA. The fraction of core promoter DNA immunoprecipitated compared to the input value was determined by real-time PCR and was expressed as percent of input (% of input) and as the fold enrichment over the fraction of *GAPDH* DNA immunoprecipitated. D. A schematic representation of the putative MafF/bZIP binding region within enhancer 2 (EnhII) of the HBV core promoter from different HBV genotypes. 10,846 HBx sequences were collected from HBVdb, The multiple sequence alignments of the MARE region were depicted by using WebLogo version 2.8. The overall height of the stack indicates the conservation at the site, while the relative frequency of each nucleic acid is shown as the height of the characters within the stack. The assays of panel B and C were performed in triplicate and include data from three independent experiments. Data are presented as mean±SD; \**p*<0.05, \*\**p*<0.01, \*\*\*\**p*<0.0001.

### 6. MafF is a competitive inhibitor of hepatocyte nuclear factor (HNF)-4α binding to HBV EnhII

HNF-4α is a transcription factor that has been previously reported to bind HBV core promoter and to induce its transcriptional activity (31, 52, 69). We found that the predicted MafF/bZIP binding region in the EnhII overlaps at it conserved 5’ region with an HNF-4α binding site that is located between nucleotides 1662 to 1674 of the HBV C_JPNAT core promoter (10) (Fig. 6A). This finding suggests the possibility that MafF may compete with HNF-4α at these binding sites within the EnhII region. To examine this possibility, we constructed a deletion mutant of EnhII/Cp (EnhII/CpΔHNF-4α#2) that extends from nt 1591 to nt 1750; this construct includes the overlapping binding regions identified for MafF and HNF-4α (i.e., HNF-4α site #1 at nt 1662–1674) but lacks the second HNF-4α binding site (HNF-4α site #2 at nt 1757–1769) as shown in Fig. 6A. We performed a ChIP assay and found that the interaction between HNF-4α and EnhII/CpΔHNF-4α #2 was significantly reduced in the presence of MafF (Fig. 6B, *p*<0.01 for % of input and *p*<0.05 for fold enrichment). Furthermore, MafF had no impact on the expression of HNF-4α (Fig. 6C). Together, these data indicated that MafF interacts directly with the HBV core promoter at the putative binding region and suppresses the transcriptional activity of the HBV core promoter by competitive inhibition of HNF-4α binding at an overlapping site in the EnhII region. This competitive suppression is due to partial overlapping of HNF4-α and the putative MafF binding regions in HBV core promoter and did not affect other host genes like ApoA1 and HNF1A known to be regulated by HNF-4α (37, 63) (Fig. 6D).

**Figure 6.**
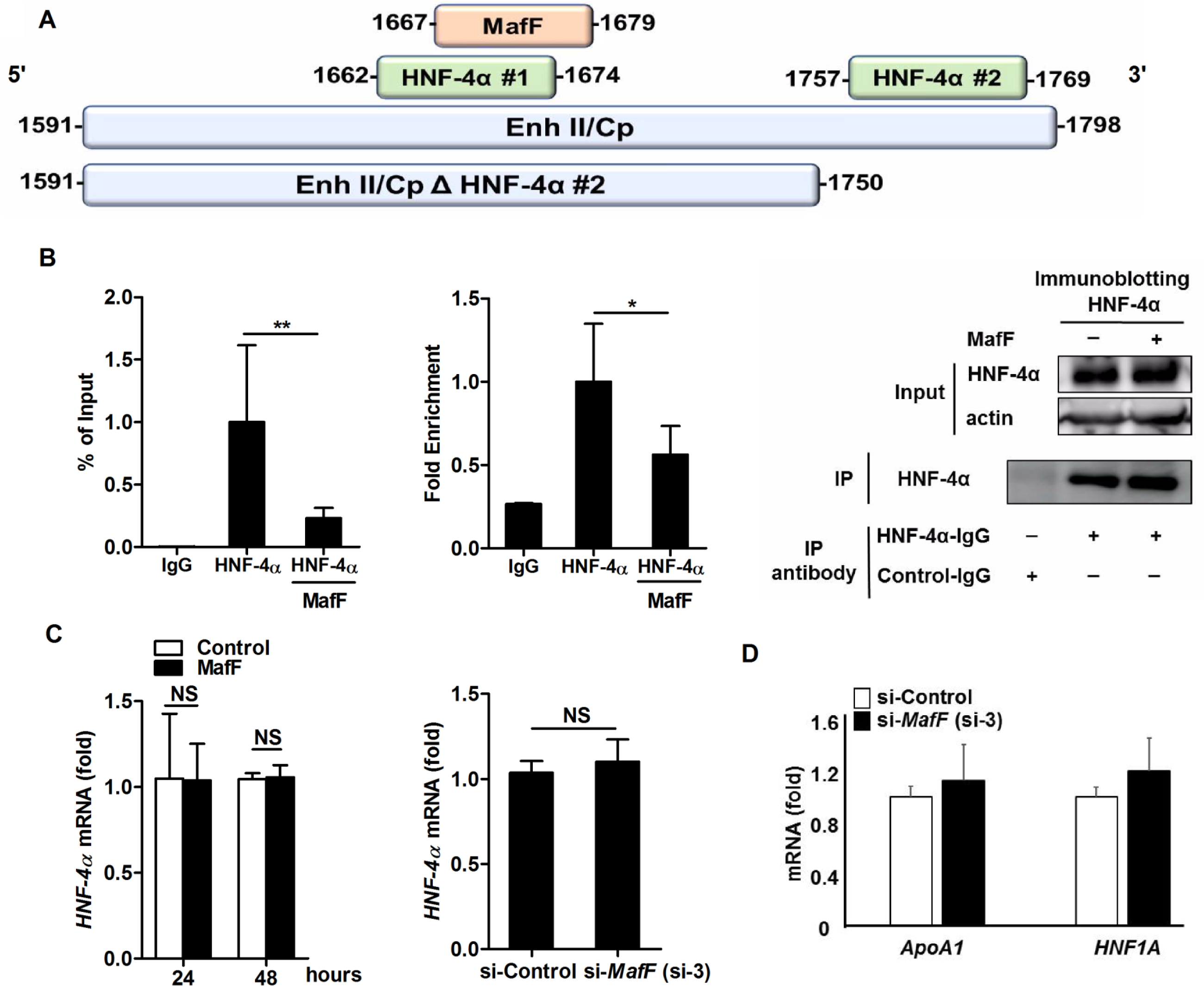
MafF competes with HNF-4α for binding to the HBV core promoter. **A.** A schematic representation of the enhancer 2 (EnhII) and the basal HBV core promoter (Cp; nt 1591-1798) featuring the putative MafF binding region (nt 1667–1679) and the two HNF-4α binding sites HNF-4α#1 (nt 1662–1674) and HNF-4α#2 (nt 1757–1769). A deletion mutant construct (EnhII/CpΔHNF-4α#2, nt 1591–1750) was prepared to eliminate HNF-4α#2. **B.** 293FT cells were co-transfected with the EnhII/CpΔHNF-4α#2-luciferase reporter plasmid, a FLAG-tagged HNF-4α expression plasmid, and a MafF (or control) expression plasmid at a ratio of 1:1:2. At two days post-transfection, cell lysates were collected and two aliquots (1/10 volume each) were removed from each sample. One aliquot was used for the detection of HNF-4α protein (Input) and actin (loading control) by immunoblotting (lower panel); the second aliquot was used for DNA extraction and detection of HBV core promoter (Input) by real-time PCR. The remaining cell lysates (each 4/10 of the original volume) were subjected to ChIP assay using isotype control antibody (rabbit IgG) or rabbit anti-HNF-4α IgG to precipitate FLAG-tagged HNF-4α. Following immunoprecipitation (IP), 1/5 volume of each IP sample was analyzed by immunoblotting to detect HNF-4α (lower panel) and each remaining IP sample was subjected to DNA extraction and real-time PCR assay for the detection of associated HBV core promoter DNA. The fraction of core promoter DNA immunoprecipitated compared to the input value was determined by real-time PCR and was expressed as percent of input (% of input) (upper left panel) and as the fold enrichment (upper right panel) over the fraction of *GAPDH* DNA immunoprecipitated. **C. Left panel:** HepG2 cells were transfected with empty vector (control) or MafF expression vector. After 24 h or 48 h, total RNA was extracted and *HNF-4α* expression was quantified by real-time RT-PCR. The data were normalized to *GAPDH* expression and are presented as fold differences relative to the control cells. **Right panel:** HepG2 cells were transfected with control or *MafF*-targeting siRNA (si-3) and *HNF-4α* expression was evaluated 48 h later as noted just above. **D.** HepG2 cells were transfected with control or *MafF*-targeting siRNA (si-3), and *HNF1A* and *ApoA-1* expression was evaluated after 48 h as previously mentioned. All assays were performed in triplicate and data are presented from three independent experiments. Data are presented as mean±SD; \**p*<0.05, \*\**p*<0.01; NS, not significant.

### 7. IL-1β and TNF-α-mediated induction of MafF expression *in vitro*

Given these findings, we speculated that MafF expression might be induced in hepatocytes in response to HBV infection. Based on a previous report of the induction of MafF by both IL-1β and TNF-α in myometrial cells (32), and the fact that both of these cytokines have been implicated in promoting host defense against HBV, we explored the possibility that MafF might be induced by one or more of these cytokines in our *in vitro* system. As shown in Fig. 7, addition of IL-1β or TNF-α resulted in significant induction of *MafF* mRNA expression in HepG2 cells (Fig. 7A, *p*<0.0001 for each cytokine); MafF protein was also detected at higher levels in HepG2 cells exposed to each of these cytokines (Fig. 7B). NF-κB is a downstream regulatory factor that is shared by the IL-1β and TNF-α signaling pathways. We found that chemical inhibition of NF-κB activity with Bay11-7082 or BMS-3455415 suppressed the induction of *MafF* expression in response to IL-1β (Fig. 7C, D; *p*<0.05 for each of these inhibitors) and to TNF-α (Fig. 7E; *p*<0.01). These findings indicate that the IL-1β and TNF-α−mediated induction of *MafF* expression in hepatocytes is regulated by NF-κB signaling. Since we showed that MafF competes with HNF-4α for its interaction with HBV core promoter (Fig. 6), we hypothesized that this effect can be enhanced by IL-1β. We performed a ChIP assay and found that 3 hours after treatment with IL-1β, the interaction between HNF-4α and EnhII/CpΔHNF-4α #2 was significantly reduced. This effect was partially reversed when *MafF* expression was silenced (Fig. 7F, *p*<0.01 for % of input and fold enrichment). These data mechanistically explain the role of MafF on the suppression of HBV core promoter activity by the inflammatory cytokine IL-1β.

**Figure 7.**
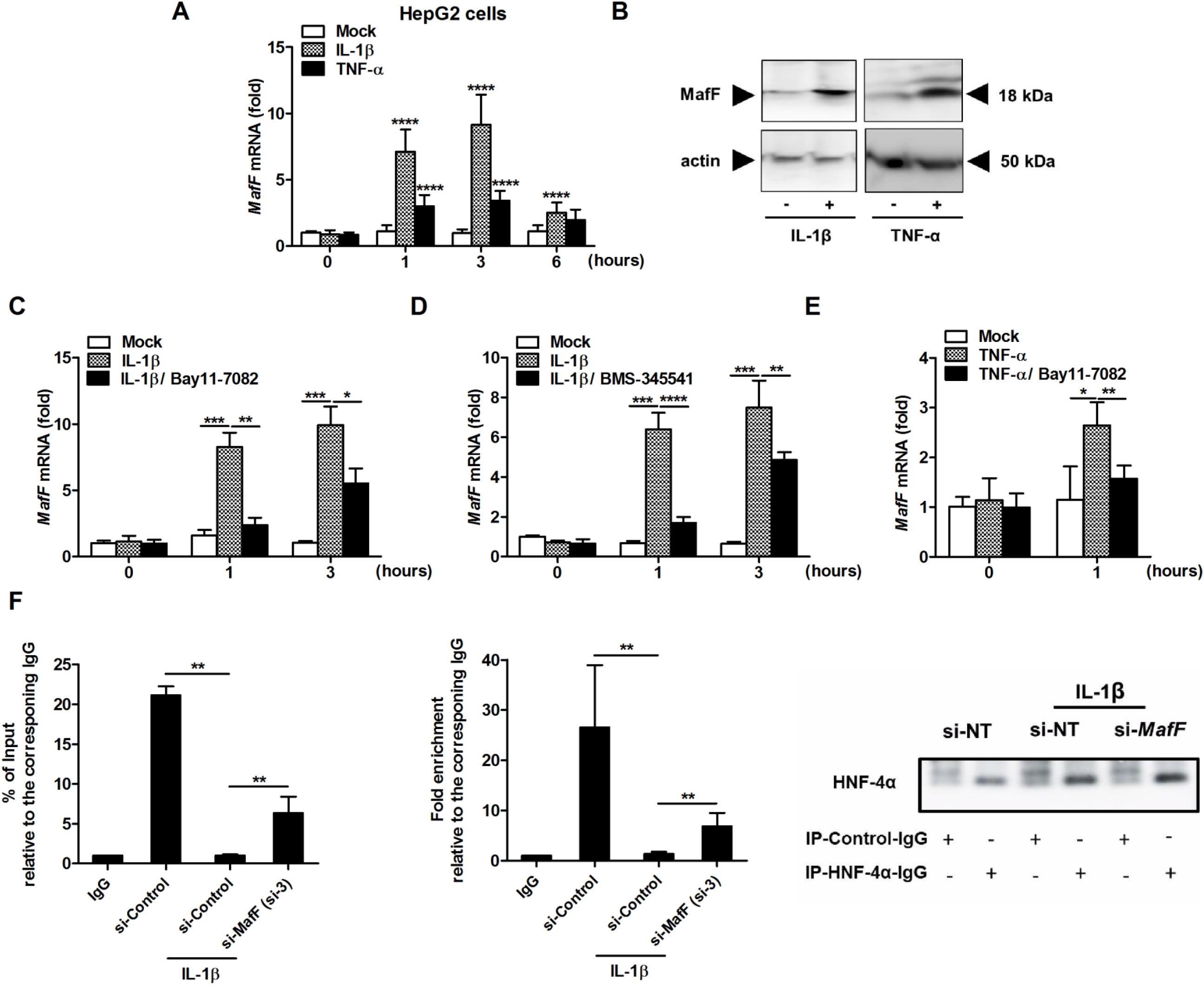
IL-1β and TNF-α induce MafF expression via NF-ҝB-mediated signaling. **A.** HepG2 cells were treated with IL-1β (1 ng/ml), TNF-α (10 ng/ml), or PBS (diluent control) for the times as indicated (hours). The cells then were lysed, total cellular RNA was extracted, and *MafF* mRNA was quantified by real-time RT-PCR. The data were normalized to the expression of *ACTB* and are shown as the fold change relative to the mean of the control group. **B.** HepG2 cells were treated for 24 h with IL-1β, TNF-α, or PBS control as in A.; the cells then were harvested and total protein was extracted. Expression of MafF (upper panel) and actin (the loading control; lower panel) was analyzed by immunoblotting. **C, D, and E.** HepG2 cells were pretreated with NF-κB inhibitors Bay11-7082, BMS-345541, or DMSO (diluent control) for 1 h and then treated with 1 ng/ml IL-1β (**C, D**), 10 ng TNF-α (**E**) or PBS (control) for 1 and 3 h. Expression of MafF was quantified by real-time RT-PCR and were normalized to the expression of the *ACTB* and shown as the fold change relative to the mean of the control group. **F.** HepG2 cells were transfected with the anti-MafF si-3 and EnhII/Cp ΔHNF-4α#2-luciferase reporter plasmid. At two days post-transfection, cells were treated with IL-1β (1 ng/mL) or mock for 3 hours, then cell lysates were collected and two aliquots (1/10 volume each) were removed from each sample. One aliquot was used for the detection of HNF-4α protein (Input) and actin (loading control) by immunoblotting (lower panel); the second aliquot was used for DNA extraction and detection of HBV core promoter (Input) by real-time PCR. The remaining cell lysates (each 4/10 of the original volume) were subjected to ChIP assay using isotype control antibody (rabbit IgG) or rabbit anti-HNF-4α IgG to precipitate endogenous HNF-4α. Following immunoprecipitation (IP), 1/5 volume of each IP sample was analyzed by immunoblotting to detect HNF-4α (lower panel) and each remaining IP sample was subjected to DNA extraction and real-time PCR assay for the detection of associated EnhII/Cp Δ HNF-4α#2 DNA. The fraction of EnhII/Cp Δ HNF-4α#2 DNA immunoprecipitated compared to the input value was determined by real-time PCR and was expressed as percent of input (% of input) (upper left panel) and as the fold enrichment (upper right panel) over the fraction of *GAPDH* DNA immunoprecipitated. All assays were performed in triplicate and including the results from three (panels A, B, and C) or two (D, E and F) independent experiments. Data are presented as mean±SD; *p<0.05, \*\**p*<0.01, \*\*\**p*<0.001, \*\*\*\**p*< 0.0001.

### 8. MafF targets HBV infection in human primary hepatocytes

Loss-of-function experiment in the primary hepatocytes is the ideal experimental platform to analyze the physiological significance of endogenous MafF on the HBV life cycle. We silenced *MafF* expression in human primary hepatocytes (PXB cells) using two independent siRNAs, including si-3, which efficiently targets the *MafF* transcript, and si-2, which was associated with a negligible silencing efficiency (Fig. 1C and Fig. 8A, upper panels) followed by infection with HBV (genotype D). *MafF* silencing in response to si-3 resulted in significant induction of HBV-pgRNA, while administration of si-2 did not yield a similar effect (Fig. 8A, lower panels; *p*<0.05). In all experiments, transcription of pgRNA was inversely associated with expression of *MafF* (Fig. 8B, *p*=0.008); these findings confirmed the role of endogenous MafF with respect to the regulation of HBV-pgRNA transcription. To confirm our earlier findings documenting induction of *MafF* by IL-1β and TNF-α, we treated PXB cells with both cytokines and observed a significant increase in *MafF* mRNA (Fig. 8C, *p*<0.05 for each cytokine).

**Figure 8.**
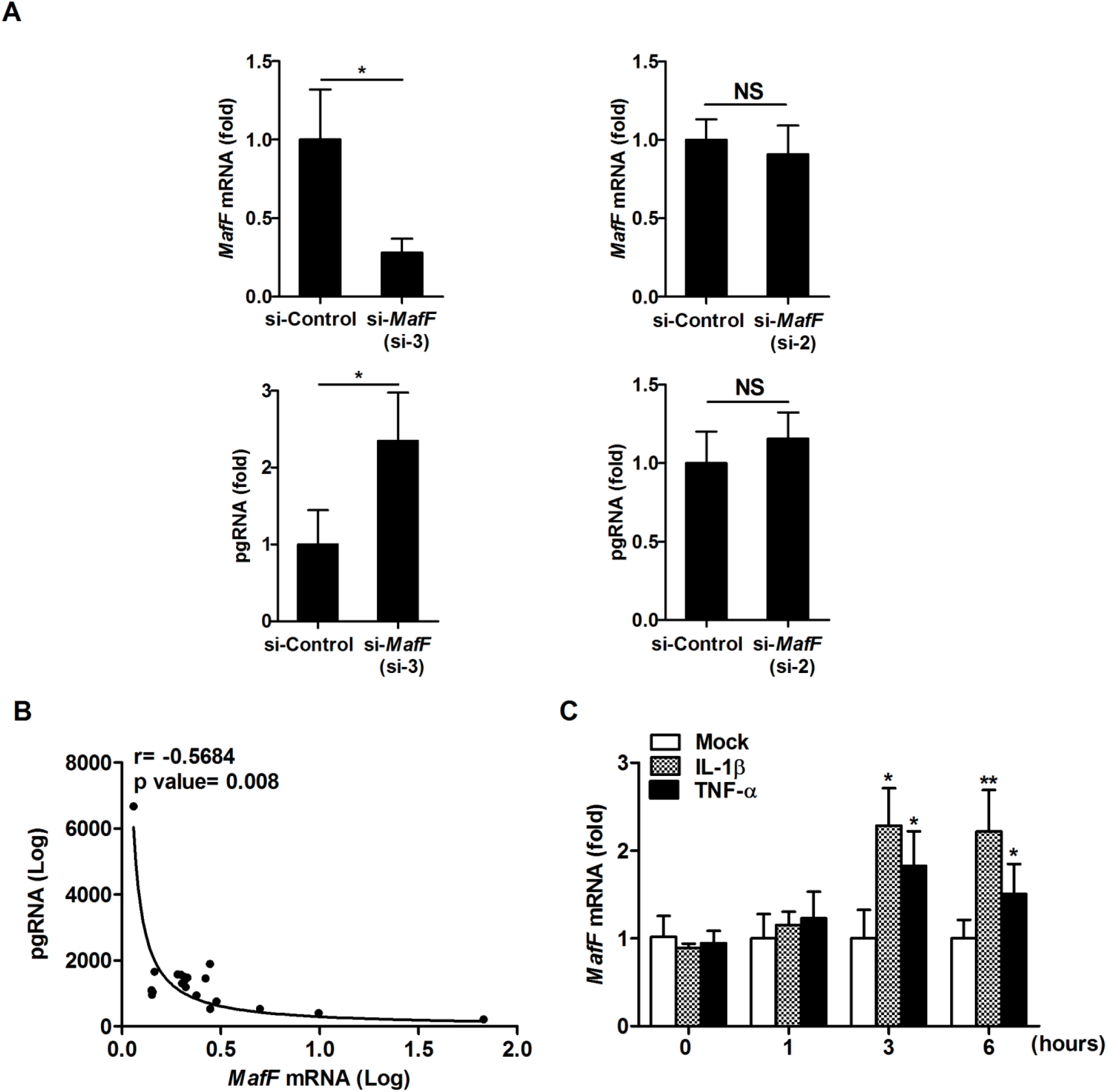
MafF suppresses HBV infection in primary human hepatocytes (PXB cells). **A.** Primary hepatocytes (PXB cells) were infected with HBV virions at 5,000 GEq per cell. After 3 days, the cells were transfected with control or *MafF*-targeting siRNAs (si-2 and si-3); at 4 days after transfection, total RNA was extracted. **Upper panel:** *MafF* expression level was quantified by real-time RT-PCR and normalized to the expression of *ACTB*. **Lower panel:** Levels of pgRNA were quantified by real-time RT-PCR using a standard curve quantification method. Data are presented as fold differences relative to the control siRNA-transfected cells. **B.** Correlation between expression of *MafF* mRNA and pgRNA in HBV-infected and siRNA-transfected PXB cells as described in A. **C.** Primary hepatocytes (PXB) cells were treated with IL-1β (at 10 ng/ml), TNF-α (at 10 ng/ml), or PBS (diluent control) for the times indicated (hours). The cells then were lysed, total cellular RNA was extracted, and *MafF* mRNA was quantified by real-time RT-PCR. The data were normalized to the expression of *ACTB* and are shown as the fold change relative to the mean of the control group. D. MafF mRNA expression detected by RNA sequencing in single primary hepatocyte (PXB) infected with HBV. All assays were performed in triplicate and include data from two independent experiments. Data are presented as mean±SD; \**p*<0.05, \*\**p*<0.01; NS, not significant

### 9. *MafF* expression is higher in HBV chronically infected patients with a positive correlation to *IL-1β* and *TNF-α* expression

To explore a role for MafF in HBV infection in human subjects, we evaluated data from an open database (71), and found that *MafF* was expressed at significantly higher levels in patients with chronic HBV compared to healthy individuals (Fig. 9A, *p*<0.0001); this was notably the case in patients undergoing immune clearance HBV (Fig. 9B, *p*<0.0001). This result confirmed the induction of *MafF* expression during active inflammation associated with this infection. This observation was strengthened by the demonstration of positive correlations between the levels of *IL-1β* and *TNF-α* transcripts and those encoding *MafF* in the immune clearance patient subset (Fig. 9C, D). Interestingly, no correlations were observed between *MafF* expression and transcripts encoding IFNs (Fig. 9E, F, G, and H). These data suggest that MafF induction associated with chronic HBV disease was unrelated to induction of IFN signaling pathways.

**Figure 9.**
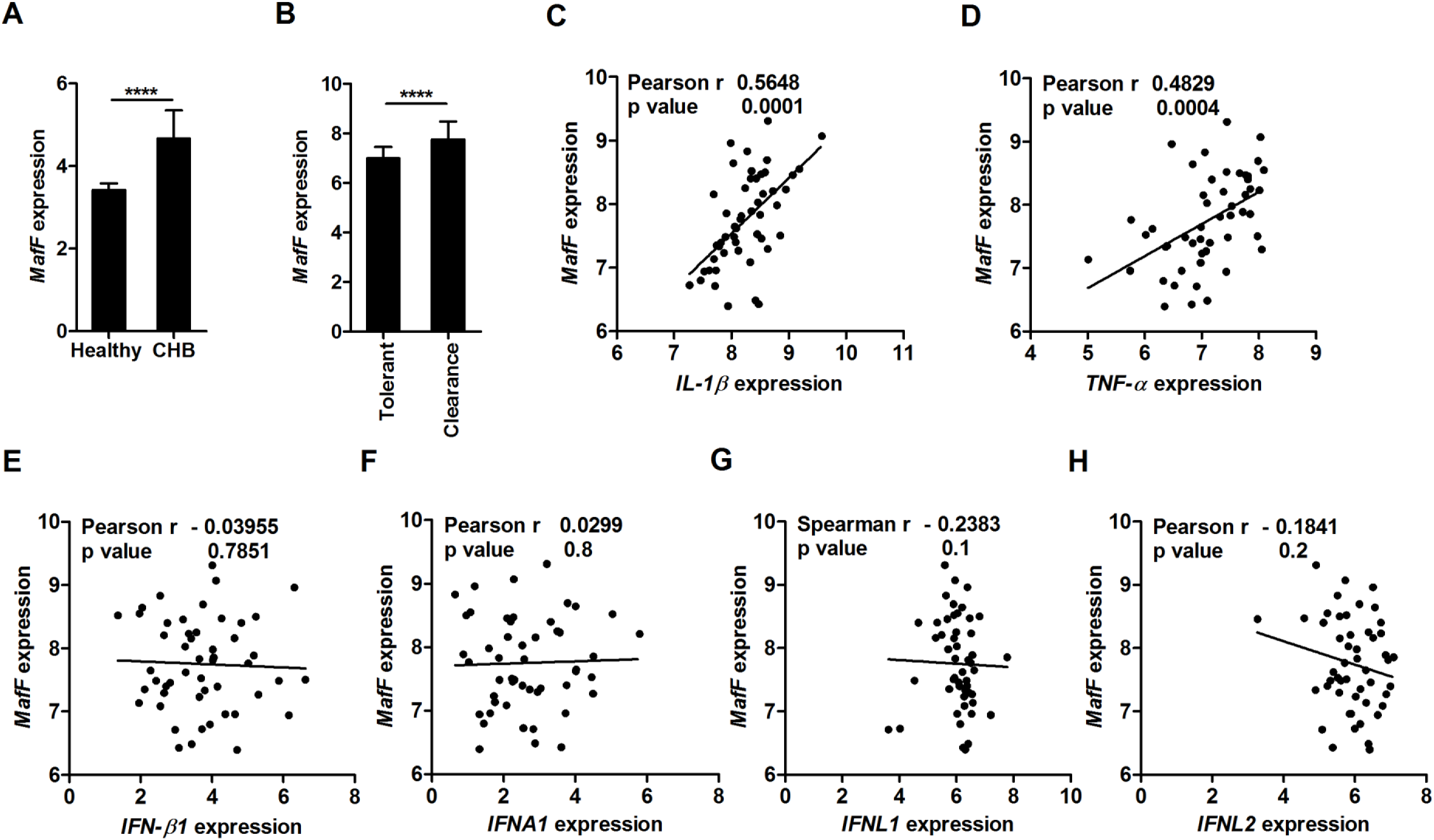
*MafF* expression is increased in patients with chronic HBV infections and is positively correlated to expression of *IL-1β* and *TNF-α* mRNAs A. *MafF* mRNA levels in the liver tissue of patients with chronic hepatitis B infection (CHB; n=122) and healthy subjects (n=6, GSE83148). B. *MafF* mRNA levels in the liver tissue of immune-tolerant (n=22) and immune clearance (n=50) HBV-infected patients (GSE65359). **C.-H.** Correlations between the expression of mRNAs encoding *MafF* and C. *IL-1β*, D. *TNF-α*, **E**. *IFN-β1*, F. *IFNA1*, **G**. *IFNL1*, and **H**. *IFNL2* in liver tissue of patients undergoing immune clearance. In panels A and D, data are presented as the mean±SD; \*\*\*\**p*<0.0001.

## Discussion

The intrinsic or innate immune response is mediated by cellular restriction factors. Many of these factors are induced by cytokines (29, 56, 58) and serve to suppress different stages of the viral life cycle, from entry to virion release (5). Several host restriction factors can suppress transcription from DNA virus promoters (57, 67). In this work, we identified MafF as a new host restriction factor that can inhibit both HBV via transcriptional suppression at targeted viral promoter. MafF significantly suppressed HBV core promoter transcription and consequently HBV-pgRNA, core protein and HBV-DNA levels. MafF-KO cells showed a significant increase of HBV core protein with no effect on HBs levels. This can be explained by the major suppressive effect exhibited by MafF on HBV-core promoter in comparison to minimal/no effect of HBV-PreS1, and PreS2 promotors respectively (Fig. 2C.)

MafF is a member of the small Maf (sMaf) family of transcription factors, a group that includes MafG (20), MafK, and MafF (9). The sMafs arebZIP-type transcription factors that bind to DNA at Maf recognition elements (MAREs). MAREs were initially defined as a 13-bp (TGCTGA(G/C)TCAGCA) or 14-bp (TGCTGA(GC/CG)TCAGCA) elements (22, 25). However, multiple studies (26, 44, 53) have presented findings suggesting heterogeneity within MARE sequences especially when sMafs heterodimerize with other bZIP-type transcription factors. Using the JASPAR database for transcription factor binding sites, we identified a sequence extending from nt 1667 to 1679 (TGGACTCTCAGCG) in the EnhII region of HBV as a potential MafF/bZIP binding region. Although we did not identify the dimerization partner of MafF in this study, we hypothesize that MafF binds to this region as a heterodimer with another bZIP transcription factor based on the weak alignment at the 5’ region of the identified MafF/bZIP binding site when compared to the palindromic MARE consensus sequence. In fact, sMafs/Bach1 dimers were previously reported to act as transcriptional repressors (60), also Bach1 binding site showed a close similarity to the identified MafF/bZIP binding sequence identified in this study (Fig. 5) highlighting the possibility that heterodimers between Bach1/MafF may be behind the MafF-mediated suppression of transcription from HBV core promoter reported in this study. Further studies need to be done to confirm this hypothesis and to identify the dimerization partner of MafF. Both ChIP and functional analysis confirmed the importance of the interaction between MafF and this specific sequence in HBV core promoter; MafF binding at this core-promoter region results in suppression of the transcriptional activity from HBV core promoter and inhibition of the HBV life cycle. Interestingly, the putative MafF/bZIP binding region in the HBV core promoter showed considerable similarity among several HBV genotypes, especially genotypes A, B, and C and ancient HBVs. Although the origin of the HBV infection in humans is still controversial; Paraskevis et al. reported that genotype C is the oldest of human HBVs (47). Indeed, by analyzing ancient HBVs derived from human skeletons or mummies of the Bronze age to the early modern age (appx. 4,500 - 250 years ago), we observed a considerable sequence similarity between the putative MafF/bZIP binding region in these old sequences and that of genotype C (47). These data suggest that MafF has continuously targeted HBV infection since the HBV infected humans (∼ 7,000 years ago). On the other hand, mutations at the 3’ region of the putative MafF/bZIP binding site are more frequent in new world HBV genotypes E, F, and G, suggesting that some HBV genotypes may acquire mutations to overcome MafF-mediated host restriction; however, whether these mutations help in evading the suppressive function of MafF and its impact on HBV life cycle still needs to be addressed.

Transcription driven from HBV core promoter is controlled by two enhancers, enhancer I (EnhI) and EnhII, the latter overlapping with the core promoter (EnhII/Cp); transcription is also modulated by a negative regulatory element (NRE) (48). Liver-enriched transcription factors, including C/EBPα, HNF-4α, HNF3, FTF/LRH-1, and HLF4, can interact with the EnhII/Cp region and thereby enhance the core promoter activity (15, 51). Negative regulation of HBV core promoter mainly takes place at the NRE, which is located immediately upstream of EnhII (38). Our analysis of the EnhII segment revealed an overlap between MafF and one of the HNF-4α binding sites located between nt 1662 to nt 1674. We identified MafF as a novel negative regulator of EnhII activity that acts via competitive inhibition of HNF-4α binding to the HBV core promoter at this site; we present this mechanism as a plausible explanation for MafF-mediated suppression of HBV infection.

The expression levels of sMafs serve as strong determinants of their overall function. An excess of sMafs may increase shift the balance toward transcriptional repression (40). As discussed previously, sMafs dimerize with CNC family proteins Nrf1, Nrf2, Nrf3, Bach1, and Bach2 (39). Furthermore, MARE consensus sites include an embedded canonical AP1 motif; as such, some Jun and Fos family factors can also heterodimerize with Maf/CNC proteins. Finally, large Maf proteins are also capable of binding at MARE elements (21, 41). Given the large number of possible homo- and heterodimeric combinations of proteins capable of binding to MAREs, transcriptional responses ranging from subtle to robust can be elicited at a single MARE site (41). Our findings revealed that MafF expression is induced by IL-1β and TNF-α in primary hepatocytes (PXBs) and that this induction was mediated by NF-κB, an inducible transcription factor that is a central regulator of immune and inflammatory responses (30). Both IL-1β and TNF-α have been associated with protection against HBV. For example, a polymorphism in the IL-1β−gene has been linked to disease progression in patients with HBV-related hepatitis (35), while TNF-α expression in hepatocytes induced by HBV (11) has been shown to decrease the extent of HBV persistence (68). We detected higher levels of *MafF* expression in patients with chronic HBV, especially among those in the immune clearance group, compared to healthy individuals. Moreover, we have also reported that IL-1β treatment significantly suppressed HNF-4α interaction with EnhII region of HBV core promoter in response to the induction of MafF expression. Correlation studies in patients’ data alone are not conclusive; however, the combination between the in vitro suppressive function of MafF and patients data suggests a possibly important role for MafF with respect to the anti-HBV effects of these cytokines in HBV-infected patients.

HBV core promoter regulates the expression of HBV precore and pgRNA transcripts. The precore-RNA serves as the template for the translation of HBV precore protein. HBV-pgRNA is translated into two proteins, HBc (the capsid-forming protein) and pol (polymerase); the HBV-pgRNA also serves as a template for HBV-DNA reverse transcription and viral replication (1). MafF inhibits HBV replication via suppressing the production of HBV-pgRNA, thereby limiting the production of the corresponding replication-associated protein (core; Fig. 3). We showed here that HBV-pgRNA titers in HBV-infected PXB were higher in cells subjected to MafF silencing; levels of HBV-pgRNA were inversely correlated with MafF mRNA levels (Fig. 7A and B). These data confirmed the importance of endogenous MafF with respect to the regulation of HBV-pgRNA transcription and viral replication. The HBV precore protein is a well-known suppressor of the anti-HBV immune response (36, 64, 70). As such, suppression of HBV precore protein expression may promote a MafF-mediated recovery of the anti-HBV immune response and enhanced viral clearance.

To summarize, the results of this work identified MafF as a novel anti-HBV. *MafF* expression was induced by both IL-1β and TNF-α in primary hepatocytes and also in patients with chronic HBV. Furthermore, MafF was shown to play an important role in the suppression of transcription from the HBV core promoter. Further analysis will be needed in order to determine whether the antiviral function of MafF is effective against other DNA viruses as well as its impact on viral evasion mechanisms.

## Materials and Methods

### Cell culture, reagents and establishment of MafF-KO cells

All the cells used in this study were maintained in culture at 37°C and 5% CO2. HepG2, HepG2-hNTCP-C4, MafF-KO HepG2-hNTCP-C4, and HepAD38.7-Tet cell lines were cultured in Dulbecco’s modified Eagle’s medium/F-12 (DMEM/F-12) GlutaMAX media (Gibco) as previously described (17). Primary human hepatocytes (Phoenixbio; PXB cells) were cultured as previously described (16). HEK 293FT cells were cultured in DMEM (Sigma) as previously described (2). For the establishment of puromycin resistant-MafF-KO cells, HepG2-hNTCP-C4 cells were co-transfected with MafF CRISPR/Cas9 KO Plasmid (h), sc-411785, and MafF HDR Plasmid (h), sc-411785-HDR, according to manufacturer instruction. Puromycin selection was conducted at 3 μg/mL. Loss of MafF expression was confirmed by immunoblotting. Myrcludex-B was kindly provided by Dr. Stephan Urban at University Hospital Heidelberg and was synthesized by CS Bio (Shanghai, China).

### Plasmid Vectors and Construction

An HBV genotype D subtype ayw replicon (62) was obtained from Addgene. HBV Ae (genotype A), HBV D_IND60, and HBV C_JPNAT are 1.24 HBV replicons, which were described previously (59). A MafF expression plasmid (pFN21AB8874) was purchased from Promega. To add a C-terminal HaloTag, the MafF-encoding sequence was subcloned into the PC14K HaloTag vector using the Carboxy Flexi system (Promega). The reporter plasmid for the HBV core promoter mutant was generated by introducing two point mutations (A1676C and C1678A) at the putative MafF binding region (Fig. 4A). Briefly, several rounds of PCR amplification were performed using pGL4.10_Ce_xmut as the template; the resulting products were digested with *Hind*III and *EcoR*I and subcloned into restriction-digested pGL4.10 (Promega). The set of primers used in the construction of the mutated core promoter include forward primers 5’-TCGAGGAATTCGGGTACTTTACCACAGGAAC-3’ and 5’-CTTGGACTCTCCGAAATGTCAACG-3’ and reverse primers 5’-TTGCCAAGCTTGAACATGAGATGATTAGGC-3’ and 5’-CGTTGACATTTCGGAGAGTCCAAG-3’. The sequence encoding HNF-4α was amplified from FR_HNF4A2 (Addgene; (61)) by PCR using primers including forward primer, 5’-AGCTAGGATCCACCATGCGACTCTCCAAAACC-3’ and reverse primer 5’-GAGTCGAATTCTTACTTGTCGTCATCGTCTTTGTAGTCAGCAACTTGCCCAAAGCG-3’. The resulting amplification product was cloned into pCDNA3.1 (Invitrogen) to yield pcDNA3.1-HNF4A-FLAG. The reporter deletion mutant EnhII/CPΔHNF-4α #2 (Fig. 5A) was constructed using pGL4.10_Ce_xmut as the template and a primer set, including forward primer 5’-TCGAGGGTACCGCCTGTAAATAGACCTATTG-3’ and reverse primer 5’-CTAACAAGCTTTCCTCCCCCAACTCCTCCC-3’; the amplification product was subcloned into pGL4.10 using *Hind*III and *Kpn*I restriction enzymes. All constructs were validated by DNA sequencing. Plasmid DNAs used in transfection experiments were purified using the Purelink Plasmid Midi Kit (Invitrogen).

### siRNA library

A Silencer Select™ Human Druggable Genome siRNA Library V4 (4397922, Thermo), was used for screening of HepG2-hNTCP-C4 cells infected with the HBV/NL reporter virus. The siRNAs were arrayed in a 96-well format; siRNAs targeting the same genes with different target sequences were distributed across three plates (A, B, and C). The following plates from this siRNA library (2200 human genes) were screened: 1-1, 1-2, 1-3, 1-4, 2-1, 2-2, 2-3, 2-4, 3-1, 5-4, 6-2, 6-3, 9-2, 11-3, 11-4, 13-3, 15-1, 15-4, 19-1, 22-2, 25-3, 25-4, 26-1, 26-2, and 26-3. Cellular viability was determined using the XTT assay (Roche) according to the manufacturer’s instructions. Wells with ≥ 20% loss of cell viability were excluded from further evaluation. Protocols for the preparation of HBV/NL and screening were as described previously (45).

### DNA and RNA transfection

Plasmid DNA transfection was performed according to the manufacturer’s guidelines, using Lipofectamine 3000 (Invitrogen) for HepG2 cells and Lipofectamine 2000 (Invitrogen) for HEK 293FT cells. Reverse siRNA transfection into HepG2-hNTCP-C4 or HepG2 was performed using Lipofectamine RNAiMAX (Thermo Fisher Scientific); forward siRNA transfection was performed in PXB cells only using Lipofectamine RNAiMAX according to the respective manufacturer’s guidelines. Silencer Select™ si-*MafF* (si-1, s24372; si-2, s24371; and si-3, s24370), si-*MafK* (s194858), si-*MafG* (s8419), and the negative control siRNA (#2) were purchased from Thermo Fisher Scientific.

### Western blot analysis

Cells were lysed with PRO-PREP protein extraction solution (Intron Biotechnology). Protein samples were separated on a 12% gel via SDS-PAGE. Immunoblotting and protein detection were performed as previously reported (2). Primary antibodies included mouse monoclonal anti-HBc (provided by Dr. Akihide Ryo, Yokohama City University), anti-Halo-tag (Promega), anti-FLAG (M2, Sigma), and anti-actin (Sigma), rabbit polyclonal anti-MafF (Protein Tech), and rabbit monoclonal anti-HNF-4α (Abcam). The band intensities were quantified by ImageJ software (NIH).

### HBV and HBV/NL preparation and infection

HBV particles carrying a chimeric HBV virus encoding NanoLuc (NL) were prepared and used as described previously (45). Briefly, HepG2 cells were transfected with a plasmid encoding HBV in which the core region was replaced by a gene encoding NL and a helper plasmid that carried an HBV genome that was defective in packaging. The resulting HBV/NL particles produced NL upon infection. HBV and HBV/NL stocks used in this study were prepared as described previously (17, 45) For infection of HepG2-hNTCP-C4 cells, the cells first were reverse-transfected with *MafF* or negative control siRNAs two days prior to the HBV infection and then infected 2 days later with inoculation of HBV or HBV/NL as described previously (17, 45); the experiment was terminated at 8 days post-infection. For PXB cells, the cells were first infected with HBV; at 3 days post-infection, the cells were transfected with the siRNAs, and the experiment was terminated at 7 days post-infection.

### RNA extraction and quantitative real-time PCR

Isolation of total cellular RNA was performed with a RNeasy Mini kit (Qiagen) according to the manufacturer’s guidelines and cDNA synthesis was performed using a Superscript VILO cDNA Synthesis Kit (Thermo Fisher Scientific). The relative levels of the *MafF* mRNA was determined using TaqMan 746 Gene Expression Assay primer-probe sets (Applied Biosystems) Hs05026540_g1, expression of *ACTB* (primer-probe set 748 Hs99999903_m1) was used as an internal control for normalization. The quantification of pgRNA, *NTCP*, *HNF-4α, APOA1, and HNF1A* was performed using Power SYBR Green PCR Master Mix (Applied Biosystems); for these transcripts, expression of *GAPDH* was used as an internal control for normalization. Data were expressed as fold change relative to the mean of the control group. The set of primers used in these assays included the Precore forward primer 5’-ACTGTTCAAGCCTCCAAGCTGT-3’ and reverse primer 5’-GAAGGCAAAAACGAGAGTAACTCCAC-3’, *NTCP* forward primer 5′-AGGGAGGAGGTGGCAATCAAGAGTGG-3’ and reverse primer 5’-CCGGCTGAAGAACATTGAGGCACTGG-3’, *HNF-4α* forward primer 5’-ACTACGGTGCCTCGAGCTGT-3’ and reverse primer 5’-GGCACTGGTTCCTCTTGTCT-3’; APOA1 forward primer 5’-CCTTGGGAAAACAGCTAAACC-3’, APOA1 reverse primer 5’-CCAGAACTCCTGGGTCACA-3’, HNF1A forward primer 5’-CCATCCTCAAAGAGCTGGAG-3’, HNF1A reverse primer 5’-TGTTGTGCTGCTGCAGGTA-3’, *GAPDH* forward primer 5’-CTTTTGCGTCGCCAG-3’ and reverse primer 5’-TTGATGGCAACAATATCCAC-3’.

### DNA extraction and cccDNA quantification

For selective extraction of cccDNAs for quantitative PCR, HBV-infected HepG2-hNTCP-C4 cells were harvested and total DNA was extracted using a Qiagen DNA extraction kit according to the manufacturer’s instructions but without the addition of Proteinase K as recommended by the concerted harmonization efforts for HBV cccDNA quantification reported in the 2019 International HBV meeting (28). Levels of cccDNA were measured by quantitative real-time PCR (qPCR) using the TaqMan Gene Expression Master Mix (Applied Biosystems), specific primers, and probe as described previously (50). Data were processed as 2^(−ΔΔCt)^ for quantification of cccDNA using chromosomal *GAPDH* DNA sequence (via primer-probe set Hs04420697_g1; Applied Biosystems) as an internal normalization control.

### Dual luciferase reporter assay

Firefly luciferase reporter plasmids carrying the entire HBV core promoter (nt 900 to 1817), Enh1/X promoter (nt 950 to 1373), preS1 promoter (nt 2707 to 2847), or preS2/S promoter (nt 2937 to 3204) were constructed as previously reported (8). HepG2 cells were co-transfected with the firefly reporter vectors and the *Renilla* luciferase plasmid pRL-TK (Promega) as an internal control. At 48 h post-transfection, the cells were lysed and luciferase activities were measured using the Dual-Luciferase Reporter Assay System (Promega). For experiments involving IL-1β, cells were treated for 3 h with IL-1β (1 ng/ml) at 48 h post-transfection followed by evaluation of dual luciferase activity.

### Quantification of HBs and HBe antigens

Cell supernatants were harvested and an ELISA was performed as described previously for HBs (65) whereas for HBe, Enzygnost HBe monoclonal, Siemens was used according to the manufacturer’s instructions.

### Indirect Immunofluorescence Analysis

Indirect immunofluorescence analysis was performed essentially as described previously (65). After fixation with 4% paraformaldehyde and permeabilization with 0.3% Triton X-100, an anti-HBc antibody (HBP-023-9, Austral Biologicals) was used as the primary antibody.

### Southern blotting assay

HepG2 cells were co-transfected with MafF-encoding or control vectors together with the HBV ayw plasmid both with or without 5 µM entecavir (Sigma) as a control. At 3 days post-transfection, core-associated DNA was isolated from intracellular viral capsids as described previously (54). Southern blot analysis to detect HBV-DNAs was performed also as described previously (2) For the detection of HBV cccDNA by southern blotting, the HBV cccDNA was extracted through protein free Hirt DNA extraction. Hirt DNA was heated at 95°C for 10 minutes to allow the denaturation of DP-rcDNA and dsDNA into single stranded DNA. The heat-treated samples were then digested with EcoRI to linearize the cccDNA. The Hirt DNA samples were then separated by agarose gel at 25 volts for 12 hours. After southern blot transfer, the DNA was hybridized with DIG Easy Hyb (Roche-11603558001) and detected with DIG wash and block buffer set (Roche-11585762001).

### Chromatin Immunoprecipitation (ChIP) assay

293FT cells were co-transfected with a MafF expression plasmid together a reporter plasmid harboring either the WT or mutated core promoter (substitution mutations in MARE) at a 4:1 ratio for assessment of the interactions between MafF and HBV core promoter. In other experiments, 293FT cells were co-transfected with plasmids encoding FLAG-tagged EnhII/CPΔHNF-4α#2, and a MafF expression plasmid (or empty vector) at a 1: 2 ratio for assessment of competitive binding of MafF and HNF-4α. Rabbit monoclonal anti-HNF-4α (Abcam, EPR16885) was used for IP. To elucidate the effect of IL-1β on MafF and HNF-4α competition, HepG2 cells were transfected with anti-MafF si-3 and plasmid encoding EnhII/CPΔHNF-4α#2. At 48 h post-transfection IL-1β was added to the culture medium (1ng/mL) for 3 hours. ChIP was carried out using a Magna ChIP G kit (Millipore) according to the manufacturer’s instructions. Anti-HNF-4α Rabbit monoclonal antibody (Abcam) was used for IP.

### Cytokine treatment and NF-**κ**B inhibitors

Responses to IL-1β and TNF-α (R&D Systems) were evaluated in HepG2 cells (at 1 ng/ml and 10 ng/ml, respectively) and in PXB cells (both at 10 ng/ml) after 1, 3, and 6 h. For the experiments involving NF-κB inhibitors, Bay11-7082 and BMS-345541 (both from Tocris) were added to final concentrations of 10 µM and 5 µM, respectively. HepG2 cells were pretreated for 1 h with each inhibitor followed by the addition of 1 ng/ml IL-1β (for 1 and 3 h) or 10 ng/ml TNF-α (for 1 h). All experiments included phosphate-buffered saline (PBS) as a diluent control for the cytokines and dimethyl sulfoxide (DMSO) as the diluent control for the NF-κB inhibitors.

### Database

Transcriptional profiling of the patients with chronic HBV (CHB) (GSE83148) and of HBV patients with immune tolerance and those undergoing HBV clearance (GSE65359) were identified in the Gene Expression Omnibus public database. Expression data for *MafF* and for genes encoding cytokines *IL-1β, TNF-α, IFNA1, IFN-β1, IFNL1, and IFNL2* were extracted by GEO2R. HBx sequences (n = 10,846) were collected from HBVdb (13), and 13 ancient HBV sequences were downloaded from the NCBI nucleotide database (18, 42, 49). For each genotype, nucleotide sequences were aligned by using MAFFT version 7.471 (23). Multiple sequence alignments of the MARE region were depicted using WebLogo version 2.8 (7). The overall height of the stack indicates the conservation at the site while the relative frequency of each nucleic acid is shown as the height of the characters within the stack.

### Statistical analysis

The data were analyzed with algorithms included in Prism (v. 5.01; GraphPad Software, San Diego, CA). Tests for normal distribution of the data were performed. Two-tailed unpaired t tests, and Mann-Whitney U tests were used for statistical analysis of parametric and non-parametric data, respectively. The correlation coefficients were determined by Pearson or Spearman correlation analysis of parametric and non-parametric data, respectively. Values of *p*≤0.05 were considered statistically significant.

## Acknowledgments

Ibrahim MK was the recipient of a JSPS postdoctoral fellowship. This study was supported by a Grant-In-Aid for JSPS Fellows (18F18098), and for Scientific Research (19K07586), and grants from the Research Program on Hepatitis from the Japan Agency for Medical Research and Development (AMED; 19fk0310104j0903, 19fk0310103j0303, 19fk0310109h0003, and 19fk0310109j0403). Myrcludex-B, a pre-S1 lipopeptide, was kindly provided by Dr. Stephan Urban at the University Hospital Heidelberg. We gratefully acknowledge Dr. Akihide Ryo (Yokohama City University School of Medicine) for the kind gift of the anti-HBV core antibody, Dr. Takayuki Murata (Fujita Health University), Dr. Iwao Kukimoto, Miss Yingfang Li, Dr. Xin Zheng, and Dr. Haruka Kudo (NIID) for productive discussion, criticism, and technical support. We are also very grateful to Dr. Haitao Guo (University of Pittsburgh) for his great support that enabled us to detect HBV-cccDNA by southern blot assay.

## References

1. Abraham, T. M., and D. D. Loeb. 2007. The topology of hepatitis B virus pregenomic RNA promotes its replication. J Virol 81:11577–84.

2. Aly, H. H., J. Suzuki, K. Watashi, K. Chayama, S. Hoshino, M. Hijikata, T. Kato, and T. Wakita. 2016. RNA Exosome Complex Regulates Stability of the Hepatitis B Virus X-mRNA Transcript in a Non-stop-mediated (NSD) RNA Quality Control Mechanism. J Biol Chem 291:15958–74.

3. Amini-Bavil-Olyaee, S., Y. J. Choi, J. H. Lee, M. Shi, I. C. Huang, M. Farzan, and J. U. Jung. 2013. The antiviral effector IFITM3 disrupts intracellular cholesterol homeostasis to block viral entry. Cell Host Microbe 13:452–64.

4. Bartenschlager, R., and H. Schaller. 1992. Hepadnaviral assembly is initiated by polymerase binding to the encapsidation signal in the viral RNA genome. EMBO J 11:3413–20.

5. Chemudupati, M., A. D. Kenney, S. Bonifati, A. Zani, T. M. McMichael, L. Wu, and J. S. Yount. 2019. From APOBEC to ZAP: Diverse mechanisms used by cellular restriction factors to inhibit virus infections. Biochim Biophys Acta Mol Cell Res 1866:382–394.

6. Chen, G., C. H. Liu, L. Zhou, and R. M. Krug. 2014. Cellular DDX21 RNA helicase inhibits influenza A virus replication but is counteracted by the viral NS1 protein. Cell Host Microbe 15:484–93.

7. Crooks, G. E., G. Hon, J. M. Chandonia, and S. E. Brenner. 2004. WebLogo: a sequence logo generator. Genome Res 14:1188–90.

8. Deng, L., X. Gan, M. Ito, M. Chen, H. H. Aly, C. Matsui, T. Abe, K. Watashi, T. Wakita, T. Suzuki, T. Okamoto, Y. Matsuura, M. Mizokami, I. Shoji, and H. Hotta. 2019. Peroxiredoxin 1, a Novel HBx-Interacting Protein, Interacts with Exosome Component 5 and Negatively Regulates Hepatitis B Virus (HBV) Propagation through Degradation of HBV RNA. J Virol 93.

9. Fujiwara, K. T., K. Kataoka, and M. Nishizawa. 1993. Two new members of the maf oncogene family, mafK and mafF, encode nuclear b-Zip proteins lacking putative trans-activator domain. Oncogene 8:2371–80.

10. Gilbert, S., L. Galarneau, A. Lamontagne, S. Roy, and L. Belanger. 2000. The hepatitis B virus core promoter is strongly activated by the liver nuclear receptor fetoprotein transcription factor or by ectopically expressed steroidogenic factor 1. J Virol 74:5032–9.

11. Gonzalez-Amaro, R., C. Garcia-Monzon, L. Garcia-Buey, R. Moreno-Otero, J. L. Alonso, E. Yague, J. P. Pivel, M. Lopez-Cabrera, E. Fernandez-Ruiz, and F. Sanchez-Madrid. 1994. Induction of tumor necrosis factor alpha production by human hepatocytes in chronic viral hepatitis. J Exp Med 179:841–8.

12. Gripon, P., I. Cannie, and S. Urban. 2005. Efficient inhibition of hepatitis B virus infection by acylated peptides derived from the large viral surface protein. J Virol 79:1613–22.

13. Hayer, J., F. Jadeau, G. Deleage, A. Kay, F. Zoulim, and C. Combet. 2013. HBVdb: a knowledge database for Hepatitis B Virus. Nucleic Acids Res 41:D566–70.

14. Hirsch, R. C., J. E. Lavine, L. J. Chang, H. E. Varmus, and D. Ganem. 1990. Polymerase gene products of hepatitis B viruses are required for genomic RNA packaging as wel as for reverse transcription. Nature 344:552–5.

15. Ishida, H., K. Ueda, K. Ohkawa, Y. Kanazawa, A. Hosui, F. Nakanishi, E. Mita, A. Kasahara, Y. Sasaki, M. Hori, and N. Hayashi. 2000. Identification of multiple transcription factors, HLF, FTF, and E4BP4, controlling hepatitis B virus enhancer II. J Virol 74:1241–51.

16. Ishida, Y., C. Yamasaki, A. Yanagi, Y. Yoshizane, K. Fujikawa, K. Watashi, H. Abe, T. Wakita, C. N. Hayes, K. Chayama, and C. Tateno. 2015. Novel robust in vitro hepatitis B virus infection model using fresh human hepatocytes isolated from humanized mice. Am J Pathol 185:1275–85.

17. Iwamoto, M., K. Watashi, S. Tsukuda, H. H. Aly, M. Fukasawa, A. Fujimoto, R. Suzuki, H. Aizaki, T. Ito, O. Koiwai, H. Kusuhara, and T. Wakita. 2014. Evaluation and identification of hepatitis B virus entry inhibitors using HepG2 cells overexpressing a membrane transporter NTCP. Biochem Biophys Res Commun 443:808–13.

18. Kahila Bar-Gal, G., M. J. Kim, A. Klein, D. H. Shin, C. S. Oh, J. W. Kim, T. H. Kim, S. B. Kim, P. R. Grant, O. Pappo, M. Spigelman, and D. Shouval. 2012. Tracing hepatitis B virus to the 16th century in a Korean mummy. Hepatology 56:1671–80.

19. Kannan, M. B., V. Solovieva, and V. Blank. 2012. The small MAF transcription factors MAFF, MAFG and MAFK: current knowledge and perspectives. Biochim Biophys Acta 1823:1841–6.

20. Kataoka, K., K. Igarashi, K. Itoh, K. T. Fujiwara, M. Noda, M. Yamamoto, and M. Nishizawa. 1995. Small Maf proteins heterodimerize with Fos and may act as competitive repressors of the NF-E2 transcription factor. Mol Cell Biol 15:2180–90.

21. Kataoka, K., M. Nishizawa, and S. Kawai. 1993. Structure-function analysis of the maf oncogene product, a member of the b-Zip protein family. Journal of Virology 67:2133–2141.

22. Kataoka, K., M. Noda, and M. Nishizawa. 1994. Maf nuclear oncoprotein recognizes sequences related to an AP-1 site and forms heterodimers with both Fos and Jun. Mol Cell Biol 14:700–12.

23. Katoh, K., and D. M. Standley. 2013. MAFFT multiple sequence alignment software version 7: improvements in performance and usability. Mol Biol Evol 30:772–80.

24. Katsuoka, F., and M. Yamamoto. 2016. Small Maf proteins (MafF, MafG, MafK): History, structure and function. Gene 586:197–205.

25. Kerppola, T. K., and T. Curran. 1994. A conserved region adjacent to the basic domain is required for recognition of an extended DNA binding site by Maf/Nrl family proteins. Oncogene 9:3149–58.

26. Kimura, M., T. Yamamoto, J. Zhang, K. Itoh, M. Kyo, T. Kamiya, H. Aburatani, F. Katsuoka, H. Kurokawa, T. Tanaka, H. Motohashi, and M. Yamamoto. 2007. Molecular basis distinguishing the DNA binding profile of Nrf2-Maf heterodimer from that of Maf homodimer. J Biol Chem 282:33681–90.

27. Krause-Kyora, B., J. Susat, F. M. Key, D. Kuhnert, E. Bosse, A. Immel, C. Rinne, S. C. Kornell, D. Yepes, S. Franzenburg, H. O. Heyne, T. Meier, S. Losch, H. Meller, S. Friederich, N. Nicklisch, K. W. Alt, S. Schreiber, A. Tholey, A. Herbig, A. Nebel, and J. Krause. 2018. Neolithic and medieval virus genomes reveal complex evolution of hepatitis B. Elife 7.

28. Lena Allweiss, M. Y., Barbara Testoni, Julie Lucifora, Chunkyu Ko, Bingqian Qu, Dieter Glebe, Elena S. kim, Mark Lutgehetmann, Stephan Urban, Guofeng Cheng, William Delaney, Massimo Levrero, Ulrike Protzer, Fabien Zoulim, Haitao Guo, Maura Dandri. 2019. Final results of the concerted harmonization efforts for HBV cccDNA quantification, 2019 International HBV Meeting, Melbourne, Australia.

29. Liang, G., G. Liu, K. Kitamura, Z. Wang, S. Chowdhury, A. M. Monjurul, K. Wakae, M. Koura, M. Shimadu, K. Kinoshita, and M. Muramatsu. 2015. TGF-beta suppression of HBV RNA through AID-dependent recruitment of an RNA exosome complex. PLoS Pathog 11:e1004780.

30. Liu, T., L. Zhang, D. Joo, and S. C. Sun. 2017. NF-kappaB signaling in inflammation. Signal Transduct Target Ther 2.

31. Lopez-Cabrera, M., J. Letovsky, K. Q. Hu, and A. Siddiqui. 1991. Transcriptional factor C/EBP binds to and transactivates the enhancer element II of the hepatitis B virus. Virology 183:825–9.

32. Massrieh, W., A. Derjuga, F. Doualla-Bell, C. Y. Ku, B. M. Sanborn, and V. Blank. 2006. Regulation of the MAFF transcription factor by proinflammatory cytokines in myometrial cells. Biol Reprod 74:699–705.

33. Meurs, E. F., Y. Watanabe, S. Kadereit, G. N. Barber, M. G. Katze, K. Chong, B. R. Williams, and A. G. Hovanessian. 1992. Constitutive expression of human double-stranded RNA-activated p68 kinase in murine cells mediates phosphorylation of eukaryotic initiation factor 2 and partial resistance to encephalomyocarditis virus growth. J Virol 66:5805–14.

34. Meyerson, N. R., L. Zhou, Y. R. Guo, C. Zhao, Y. J. Tao, R. M. Krug, and S. L. Sawyer. 2017. Nuclear TRIM25 Specifically Targets Influenza Virus Ribonucleoproteins to Block the Onset of RNA Chain Elongation. Cell Host Microbe 22:627–638 e7.

35. Migita, K., Y. Maeda, S. Abiru, M. Nakamura, A. Komori, S. Miyazoe, K. Nakao, H. Yatsuhashi, K. Eguchi, and H. Ishibashi. 2007. Polymorphisms of interleukin-1beta in Japanese patients with hepatitis B virus infection. J Hepatol 46:381–6.

36. Mitra, B., J. Wang, E. S. Kim, R. Mao, M. Dong, Y. Liu, J. Zhang, and H. Guo. 2019. Hepatitis B Virus Precore Protein p22 Inhibits Alpha Interferon Signaling by Blocking STAT Nuclear Translocation. J Virol 93.

37. Mogilenko, D. A., E. B. Dizhe, V. S. Shavva, I. A. Lapikov, S. V. Orlov, and A. P. Perevozchikov. 2009. Role of the nuclear receptors HNF4 alpha, PPAR alpha, and LXRs in the TNF alpha-mediated inhibition of human apolipoprotein A-I gene expression in HepG2 cells. Biochemistry 48:11950–60.

38. Moolla, N., M. Kew, and P. Arbuthnot. 2002. Regulatory elements of hepatitis B virus transcription. J Viral Hepat 9:323–31.

39. Motohashi, H., F. Katsuoka, J. A. Shavit, J. D. Engel, and M. Yamamoto. 2000. Positive or negative MARE-dependent transcriptional regulation is determined by the abundance of small Maf proteins. Cell 103:865–75.

40. Motohashi, H., T. O’Connor, F. Katsuoka, J. D. Engel, and M. Yamamoto. 2002. Integration and diversity of the regulatory network composed of Maf and CNC families of transcription factors. Gene 294:1–12.

41. Motohashi, H., J. A. Shavit, K. Igarashi, M. Yamamoto, and J. D. Engel. 1997. The world according to Maf. Nucleic Acids Res 25:2953–59.

42. Muhlemann, B., T. C. Jones, P. B. Damgaard, M. E. Allentoft, I. Shevnina, A. Logvin, E. Usmanova, I. P. Panyushkina, B. Boldgiv, T. Bazartseren, K. Tashbaeva, V. Merz, N. Lau, V. Smrcka, D. Voyakin, E. Kitov, A. Epimakhov, D. Pokutta, M. Vicze, T. D. Price, V. Moiseyev, A. J. Hansen, L. Orlando, S. Rasmussen, M. Sikora, L. Vinner, A. Osterhaus, D. J. Smith, D. Glebe, R. A. M. Fouchier, C. Drosten, K. G. Sjogren, K. Kristiansen, and E. Willerslev. 2018. Ancient hepatitis B viruses from the Bronze Age to the Medieval period. Nature 557:418–423.

43. Nassal, M. 2015. HBV cccDNA: viral persistence reservoir and key obstacle for a cure of chronic hepatitis B. Gut 64:1972–84.

44. Ney, P. A., B. P. Sorrentino, C. H. Lowrey, and A. W. Nienhuis. 1990. Inducibility of the HS II enhancer depends on binding of an erythroid specific nuclear protein. Nucleic Acids Res 18:6011–7.

45. Nishitsuji, H., S. Ujino, Y. Shimizu, K. Harada, J. Zhang, M. Sugiyama, M. Mizokami, and K. Shimotohno. 2015. Novel reporter system to monitor early stages of the hepatitis B virus life cycle. Cancer Sci 106:1616–24.

46. Orzalli, M. H., and D. M. Knipe. 2014. Cellular sensing of viral DNA and viral evasion mechanisms. Annu Rev Microbiol 68:477–92.

47. Paraskevis, D., G. Magiorkinis, E. Magiorkinis, S. Y. Ho, R. Belshaw, J. P. Allain, and A. Hatzakis. 2013. Dating the origin and dispersal of hepatitis B virus infection in humans and primates. Hepatology 57:908–16.

48. Park, Y. K., E. S. Park, D. H. Kim, S. H. Ahn, S. H. Park, A. R. Lee, S. Park, H. S. Kang, J. H. Lee, J. M. Kim, S. K. Lee, K. H. Lim, N. Isorce, S. Tong, F. Zoulim, and K. H. Kim. 2016. Cleaved c-FLIP mediates the antiviral effect of TNF-alpha against hepatitis B virus by dysregulating hepatocyte nuclear factors. J Hepatol 64:268–277.

49. Patterson Ross, Z., J. Klunk, G. Fornaciari, V. Giuffra, S. Duchene, A. T. Duggan, D. Poinar, M. W. Douglas, J. S. Eden, E. C. Holmes, and H. N. Poinar. 2018. The paradox of HBV evolution as revealed from a 16th century mummy. PLoS Pathog 14:e1006750.

50. Qu, B., Y. Ni, F. A. Lempp, F. W. R. Vondran, and S. Urban. 2018. T5 Exonuclease Hydrolysis of Hepatitis B Virus Replicative Intermediates Allows Reliable Quantification and Fast Drug Efficacy Testing of Covalently Closed Circular DNA by PCR. J Virol 92.

51. Quarleri, J. 2014. Core promoter: a critical region where the hepatitis B virus makes decisions. World J Gastroenterol 20:425–35.

52. Raney, A. K., J. L. Johnson, C. N. Palmer, and A. McLachlan. 1997. Members of the nuclear receptor superfamily regulate transcription from the hepatitis B virus nucleocapsid promoter. J Virol 71:1058–71.

53. Rushmore, T. H., M. R. Morton, and C. B. Pickett. 1991. The antioxidant responsive element. Activation by oxidative stress and identification of the DNA consensus sequence required for functional activity. J Biol Chem 266:11632–9.

54. Sakurai, F., S. Mitani, T. Yamamoto, K. Takayama, M. Tachibana, K. Watashi, T. Wakita, S. Iijima, Y. Tanaka, and H. Mizuguchi. 2017. Human induced-pluripotent stem cell-derived hepatocyte-like cells as an in vitro model of human hepatitis B virus infection. Sci Rep 7:45698.

55. Sandelin, A., W. Alkema, P. Engstrom, W. W. Wasserman, and B. Lenhard. 2004. JASPAR: an open-access database for eukaryotic transcription factor binding profiles. Nucleic Acids Res 32:D91–4.

56. S. Schoggins, J. W., D. A. MacDuff, N. Imanaka, M. D. Gainey, B. Shrestha, J. L. Eitson, K. B. Mar, R. B. Richardson, A. V. Ratushny, V. Litvak, R. Dabelic, B. Manicassamy, J. D. Aitchison, Aderem, R. M. Elliott, A. Garcia-Sastre, V. Racaniello, E. J. Snijder, W. M. Yokoyama, M. Diamond, H. W. Virgin, and C. M. Rice. 2014. Pan-viral specificity of IFN-induced genes reveals new roles for cGAS in innate immunity. Nature 505:691–5.

57. Schreiner, S., S. Kinkley, C. Burck, A. Mund, P. Wimmer, T. Schubert, P. Groitl, H. Will, and T. Dobner. 2013. SPOC1-mediated antiviral host cell response is antagonized early in human adenovirus type 5 infection. PLoS Pathog 9:e1003775.

58. Shiromoto, F., H. H. Aly, H. Kudo, K. Watashi, A. Murayama, N. Watanabe, X. Zheng, T. Kato, K. Chayama, M. Muramatsu, and T. Wakita. 2018. IL-1beta/ATF3-mediated induction of Ski2 expression enhances hepatitis B virus x mRNA degradation. Biochem Biophys Res Commun 503:1854–1860.

59. Sugiyama, M., Y. Tanaka, T. Kato, E. Orito, K. Ito, S. K. Acharya, R. G. Gish, A. Kramvis, T. Shimada, N. Izumi, M. Kaito, Y. Miyakawa, and M. Mizokami. 2006. Influence of hepatitis B virus genotypes on the intra- and extracellular expression of viral DNA and antigens. Hepatology 44:915–24.

60. Sun, J., A. Muto, H. Hoshino, A. Kobayashi, S. Nishimura, M. Yamamoto, N. Hayashi, E. Ito, and K. Igarashi. 2001. The promoter of mouse transcription repressor bach1 is regulated by Sp1 and trans-activated by Bach1. J Biochem 130:385–92.

61. Thomas, H., S. Senkel, S. Erdmann, T. Arndt, G. Turan, L. Klein-Hitpass, and G. U. Ryffel. 2004. Pattern of genes influenced by conditional expression of the transcription factors HNF6, HNF4alpha and HNF1beta in a pancreatic beta-cell line. Nucleic Acids Res 32:e150.

62. Wang, H., S. Kim, and W. S. Ryu. 2009. DDX3 DEAD-Box RNA helicase inhibits hepatitis B virus reverse transcription by incorporation into nucleocapsids. J Virol 83:5815–24.

63. Wang, H., P. Maechler, P. A. Antinozzi, K. A. Hagenfeldt, and C. B. Wollheim. 2000. Hepatocyte nuclear factor 4alpha regulates the expression of pancreatic beta -cell genes implicated in glucose metabolism and nutrient-induced insulin secretion. J Biol Chem 275:35953–9.

64. Wang, Y., L. Cui, G. Yang, J. Zhan, L. Guo, Y. Chen, C. Fan, D. Liu, and D. Guo. 2019. Hepatitis B e Antigen Inhibits NF-kappaB Activity by Interrupting K63-Linked Ubiquitination of NEMO. J Virol 93.

65. Watashi, K., G. Liang, M. Iwamoto, H. Marusawa, N. Uchida, T. Daito, K. Kitamura, M. Muramatsu, H. Ohashi, T. Kiyohara, R. Suzuki, J. Li, S. Tong, Y. Tanaka, K. Murata, H. Aizaki, and T. Wakita. 2013. Interleukin-1 and tumor necrosis factor-alpha trigger restriction of hepatitis B virus infection via a cytidine deaminase activation-induced cytidine deaminase (AID). J Biol Chem 288:31715–27.

66. Wieland, S., R. Thimme, R. H. Purcell, and F. V. Chisari. 2004. Genomic analysis of the host response to hepatitis B virus infection. Proc Natl Acad Sci U S A 101:6669–74.

67. Wustenhagen, E., F. Boukhallouk, I. Negwer, K. Rajalingam, F. Stubenrauch, and L. Florin. 2018. The Myb-related protein MYPOP is a novel intrinsic host restriction factor of oncogenic human papillomaviruses. Oncogene 37:6275–6284.

68. Xia, Y., D. Stadler, J. Lucifora, F. Reisinger, D. Webb, M. Hosel, T. Michler, K. Wisskirchen, X. Cheng, K. Zhang, W. M. Chou, J. M. Wettengel, A. Malo, F. Bohne, D. Hoffmann, F. Eyer, R. Thimme, C. S. Falk, W. E. Thasler, M. Heikenwalder, and U. Protzer. 2016. Interferon-gamma and Tumor Necrosis Factor-alpha Produced by T Cells Reduce the HBV Persistence Form, cccDNA, Without Cytolysis. Gastroenterology 150:194–205.

69. Yu, X., and J. E. Mertz. 1997. Differential regulation of the pre-C and pregenomic promoters of human hepatitis B virus by members of the nuclear receptor superfamily. J Virol 71:9366–74.

70. Yu, Y., P. Wan, Y. Cao, W. Zhang, J. Chen, L. Tan, Y. Wang, Z. Sun, Q. Zhang, Y. Wan, Y. Zhu, F. Liu, K. Wu, Y. Liu, and J. Wu. 2017. Hepatitis B Virus e Antigen Activates the Suppressor of Cytokine Signaling 2 to Repress Interferon Action. Sci Rep 7:1729.

71. Zhou, W., Y. Ma, J. Zhang, J. Hu, M. Zhang, Y. Wang, Y. Li, L. Wu, Y. Pan, Y. Zhang, X. Zhang, Z. Zhang, H. Li, L. Lu, L. Jin, J. Wang, Z. Yuan, and J. Liu. 2017. Predictive model for inflammation grades of chronic hepatitis B: Large-scale analysis of clinical parameters and gene expressions. Liver Int 37:1632–1641.

